# HARMONY: A large-scale harmonized neuroimaging dataset for research on anxious misery disorders

**DOI:** 10.64898/2026.06.26.732748

**Authors:** Setthanan Jarukasemkit, Michael P. Harms, Petra Lenzini, Andrew Chen, Matthew F. Glasser, Kassandra Hamilton, Liangfang Li, Xiaoke Luo, Michael Myers, Adam R. Pines, Erin Reid, Leonardo Tozzi, Artemis Zavaliangos-Petropulu, Jiahe Zhang, Susan Whitfield-Gabrieli, Katherine L. Narr, Leanne M. Williams, Yvette Sheline, Janine D. Bijsterbosch

## Abstract

Patterns of brain circuit dysfunction underlying depression and anxiety have been increasingly characterized, including dimensional and subtype variation. A key challenge is determining how such patterns generalize across populations and measurement frameworks. Here, we introduce HARMONY, a harmonized multimodal neuroimaging dataset supporting large-scale investigation of brain–behavior associations across symptom-defined dimensions. HARMONY integrates four Human Connectome Project–style Connectomes Related to Human Disease cohorts spanning adolescence to later adulthood and capturing anxious misery symptoms. The resource combines standardized HCP-style preprocessing, quality control, imaging-derived phenotypes, and harmonized symptom measures into a clinically enriched public dataset. Proof-of-concept analyses using HARMONY showed that pooling heterogeneous cohorts increased statistical power for detecting associations between imaging-derived phenotypes and anhedonia and depression severity. Effect sizes remained modest, consistent with symptom-based measures across heterogeneous samples. Functional imaging-derived phenotypes showed the strongest multivariate predictive performance. In summary, HARMONY provides a large multi-cohort resource for reproducible mental health neuroimaging research.

## Introduction

Since the introduction of neuroimaging into psychiatry, alterations in brain structure and function have been linked to symptoms of anxious misery, a construct referring to the overlap between depression and anxiety-related symptoms^1–5^. Landmark early findings using magnetic resonance imaging shifted depression research from a primary focus on neurochemistry toward neuroanatomy and distributed brain networks with implications for treatment^6–8^. Subsequent key findings have linked clinical depression to altered brain structure and function that change with treatment^9,10^. Complementary key findings have demonstrated that the heterogeneity of depression and anxiety forms reproducible subtypes based on functional brain circuits^11,12^. In younger individuals, landmark functional MRI (fMRI) connectivity findings highlight the potential to predict future onset of depression and allied disorders^13^. Complementary emerging approaches have integrated resting fMRI data across samples highlighting the power of large consortia datasets^14^ and the pooling of structural MRI (sMRI) across multiple datasets^15^. Parallel cohort efforts have focused on using structural imaging to refine the diagnosis of major depression versus healthy controls^16^.

Depression and anxiety disorders are highly heterogeneous, with individuals presenting similar symptom profiles despite fundamentally different underlying neurobiological mechanisms^17,18^. The need to disentangle this neural heterogeneity was recognized in the National Institute of Mental Health (NIMH)’s Research Domain Criteria (RDoC) framework, which positioned the field to use a neurobiologically based framework in psychiatry^19^. Advances in neuroimaging have identified reproducible patterns of brain circuit dysfunction that capture this heterogeneity and move beyond symptom-based definitions alone^17,18^. However, translating these insights into broadly generalizable frameworks remains a key challenge, particularly when attempting to characterize variation across populations using diverse datasets and measurement approaches.

The Human Connectome Project (HCP) launched in 2012^20^, was a seminal initiative using MRI to map the human brain at scale. HCP introduced accelerated multiband sequences for functional and diffusion MRI, alongside high-resolution structural imaging^21–23^. The HCP-Young Adult (HCP-YA) provided high-quality multimodal neuroimaging for 1200 healthy individuals, and enabled the establishment of connectome-specific processing pipelines and a high resolution ‘grayordinate’-based imaging format that captures data from surface-based processing^24^. Reflecting the vision of HCP, multiple connectome projects of healthy individuals encompassing the full lifespan were completed^25–27^. These cohorts provided a foundation for the launch of Connectomes Related to Human Disease (CRHD) projects in 14 disease-specific cohorts across a diverse set of brain disorders. Four CRHD projects, led by authors SWG, LMW, YS, and KN (Fig. 1^28^), focused on depression and anxiety: Connectomes Related to Anxiety and Depression in Adolescents (HCP-ADA, also called BANDA; PI Whitfield-Gabrieli^29,30^), Human Connectome Project for Disordered Emotional States (HCP-DES, also called STACT; PI Williams^31^), Dimensional Connectomics of Anxious Misery (HCP-DAM, also called ANXPE; PI Sheline^32^), and Perturbation of the Treatment Resistant Depression Connectome by Fast-acting Therapies (HCP-PDC, also called MDD; PI Narr). To align with NIMH Data Archive (NDA) naming conventions, we refer to these cohorts as HCP-BANDA, HCP-DES, HCP-DAM, and HCP-PDC throughout this manuscript. The goal of the HARMONY data resource is to combine these four CRHD cohorts related to anxious misery into a harmonized dataset for broader scale population level analyses.

**Figure 1:**
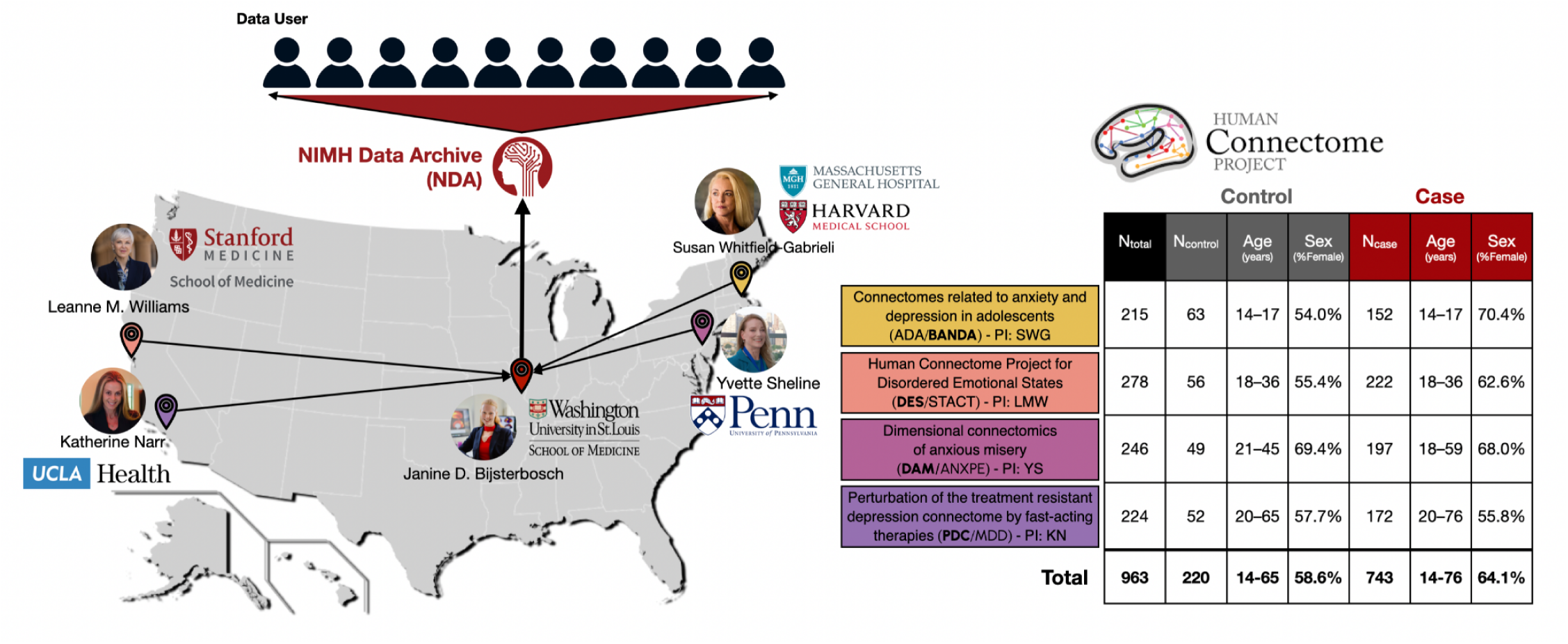
Overview of HARMONY dataset which represents the combination of four source Connectomes Related to Human Disease (CRHD) projects led by authors SWG, LMW, YS, and KN: Connectomes Related to Anxiety and Depression in Adolescents (HCP-ADA, also called BANDA; PI Whitfield-Gabrieli, Northeastern University;^29,30^), Human Connectome Project for Disordered Emotional States, Stanford University (HCP-DES, also called STACT; PI Williams;^31^), Dimensional Connectomics of Anxious Misery, University of Pennsylvania (HCP-DAM, also called ANXPE; PI Sheline^32^), and Perturbation of the Treatment Resistant Depression Connectome by Fast-acting Therapies, UCLA (HCP-PDC, also called MDD; PI Narr). Alternative cohort names are shown based on prior literature, and the bolded names in the figure are used consistently throughout the manuscript. Building on prior efforts developed by the Connectome Coordination Facility^33^, Washington University in Saint Louis served as the data collation and processing center. Each dataset acquired multimodal neuroimaging data on (N=152-222) cases meeting criteria for anxious misery (see Methods) and on (N=49-63) healthy controls. See supplementary Table S1 and Fig. S1 for a detailed breakdown of symptom severity across CRHD cohorts and supplementary Table S2 for detailed neuroimaging acquisition protocols for each cohort.

All four CRHD projects focused on anxious misery symptoms, each including approximately 150–220 cases (Fig. 1). Clinically, the cohorts were designed around distinct developmental stages, symptom severity, and treatment aspects. Specifically, HCP-BANDA included adolescents with anxiety and/or depressive disorders; HCP-DES aimed to recruit treatment-free young adults with significant anhedonia, anxious arousal, concentration, rumination, or tension symptoms; and HCP-DAM used a normative approach inclusion criterion based on Neuroticism scores ≥1 standard deviation above the control mean. In contrast, HCP-PDC is a treatment cohort focused on treatment resistant depression, including participants who had failed to respond to at least two antidepressants, were in a current depressive episode lasting at least 6 months, and had not received neuromodulation or ketamine within the previous 6 months^28^. Therefore, these four CRHD cohorts expectedly differ along the clinical spectrum of anxious misery, while collectively providing broad and complementary coverage (see Supplementary Fig. S1 for details).

Combined and harmonized datasets are particularly valuable for enabling population-level analyses that map imaging features to broad symptom constructs across cohorts^43^. Efforts such as the YaleNeuroConnect (N=302) and the Transdiagnostic Connectome Project (N=241) datasets, and the ENIGMA MDD and COORDINATE MDD consortias reflect this direction^44–47^. Complementary efforts focused on standardized imaging of specific neural systems and defined symptoms across decades of treatment response share a related focus on accelerating reproducible imaging for clinically translatable precision medicine in mental health^48^. Building on prior federal and investigator investment in CRHD data acquisition, there is a clear opportunity to aggregate the four CRHD datasets on anxious misery into a rigorously processed resource designed for cross-cohort, data-driven association analyses.

The CRHD initiative developed common strategies for acquiring and sharing connectome neuroimaging data, but combining CRHD datasets still presents some challenges^49^. In addition to well-recognized site and scanner effects^50,51^, the four cohorts differ systematically on dimensions reflecting their original scientific objectives: symptom severity (highest in HCP-PDC), age range (youngest in HCP-BANDA), and the inclusion of naturalistic follow-up (HCP-BANDA, HCP-DES, HCP-DAM) or specific treatment interventions (HCP-PDC). Further heterogeneity arises from the use of partially different instruments to assess symptom severity across cohorts. Beyond these differences related to study designs and source data acquisition, practical barriers to integration remain because public distribution through the National Institute of Mental Health Data Archive (NDA) primarily provides access to unprocessed imaging data and only cohort-specific metadata. Therefore, secondary data analyses would be facilitated by sharing imaging data pre-processed with uniform pipelines, aggregate symptom and other meta-data, and harmonized imaging summary measures. Therefore, we developed the HARMONY dataset by implementing three key steps: (i) standardized HCP-style preprocessing with rigorous quality control, (ii) collation of symptom measures across cohorts, and (iii) extraction and harmonization of imaging-derived phenotypes (IDPs)

Imaging-derived phenotypes have become a practical framework in large-scale neuroimaging resources such as the UK Biobank^52^. IDPs span from expert-curated summary measures to data-driven multimodal phenotypes. In HARMONY, we focused on widely used, interpretable IDPs with established relevance to mental health research across three imaging modalities and multiple atlases. Specifically, our IDPs cover regional morphology^53^, cortical thickness^54^, and myelin^55^ for structural MRI (sMRI); functional amplitude^56^ and connectivity^57^ for resting-state functional MRI (rsfMRI); and diffusion-tensor white matter microstructure^58^ together with neurite-related microstructure^59^ for diffusion MRI (dMRI).

Prior multi-site neuroimaging studies have demonstrated the importance of post-acquisition harmonization to mitigate batch effects, which are scanner- and site-related variation that can confound the results and limit reproducibility in downstream analyses^60^. Among retrospective harmonization approaches, ComBat-based methods are widely used to adjust for site-related differences in mean and variance^51,61^. Extensions such as CovBat further account for covariance structure^62^, which is particularly relevant for multivariate analyses of IDPs. Incorporating generalized additive models (GAM) within this framework (e.g., ComBat-GAM and CovBat-GAM) additionally allows for flexible modeling of nonlinear biological effects, such as age, helping preserve meaningful brain–behavior associations across harmonized datasets^63^. To preserve symptom-related variance across sites, we aimed to derive the harmonization model using only the controls from each site.

In summary, HARMONY provides a high quality HCP-style harmonized neuroimaging resource with standardized preprocessing and quality-control metrics, enabling immediate use for cross-cohort analyses of anxious misery constructs. Across the integrated dataset, the pooled cohorts offer increased statistical power for detecting subtle associations between broad brain-symptom associations, while post-acquisition harmonization procedures reduced site-related variance across all released IDPs. In proof-of-concept analyses, we observed that functional IDPs showed the strongest multivariate predictions, highlighting their utility for population-level association analyses of symptom-related variation.

## Results

HARMONY offers a shared data resource for studying neural dimensions of anxious misery. The results are split into four sections. In section 1, we present the harmonized HCP-style preprocessing pipeline that was applied to all data and report information regarding imaging data quality. In section 2, we cover the extraction of summary-level tabular IDPs from structural, diffusion, and resting state functional MRI modalities and their further harmonization (after pipeline processing). In section 3 we introduce proof-of-concept multivariate and univariate brain wide association results to demonstrate the value of the HARMONY resource for neuroimaging research in anxious misery. Lastly, in section 4 we share details on where to access the HARMONY data resource and guidance for users.

### Preprocessing and quality control of neuroimaging data

All data were identically preprocessed on the same servers at Washington University in St Louis. An updated version of the HCP preprocessing pipeline was implemented. Briefly, this pipeline included the published minimal HCP preprocessing pipeline^24^ implemented in QuNex^64,65^ with two key extensions: (i) rigorous quality control of surface registration and (ii) extended fMRI preprocessing pipeline (Fig. 2). Specifically, all Freesurfer cortical ribbon extractions were manually reviewed by author ER and edited where needed (in 5.3% of participants; see Supplementary Table S3 for details). In these subjects, all subsequent preprocessing steps were repeated following manual editing of the surface. Extended fMRI preprocessing included ICA-FIX cleanup to remove spatially structured noise components^66,67^, cross-participant surface registration using multimodal surface matching (‘MSMAll’) to improve functional alignment^68,69^, ‘Reclean’ module designed to improve upon ICA-FIX classification of signal and noise components^70^ and temporal ICA cleanup to remove unstructured noise components^71^.

**Figure 2:**
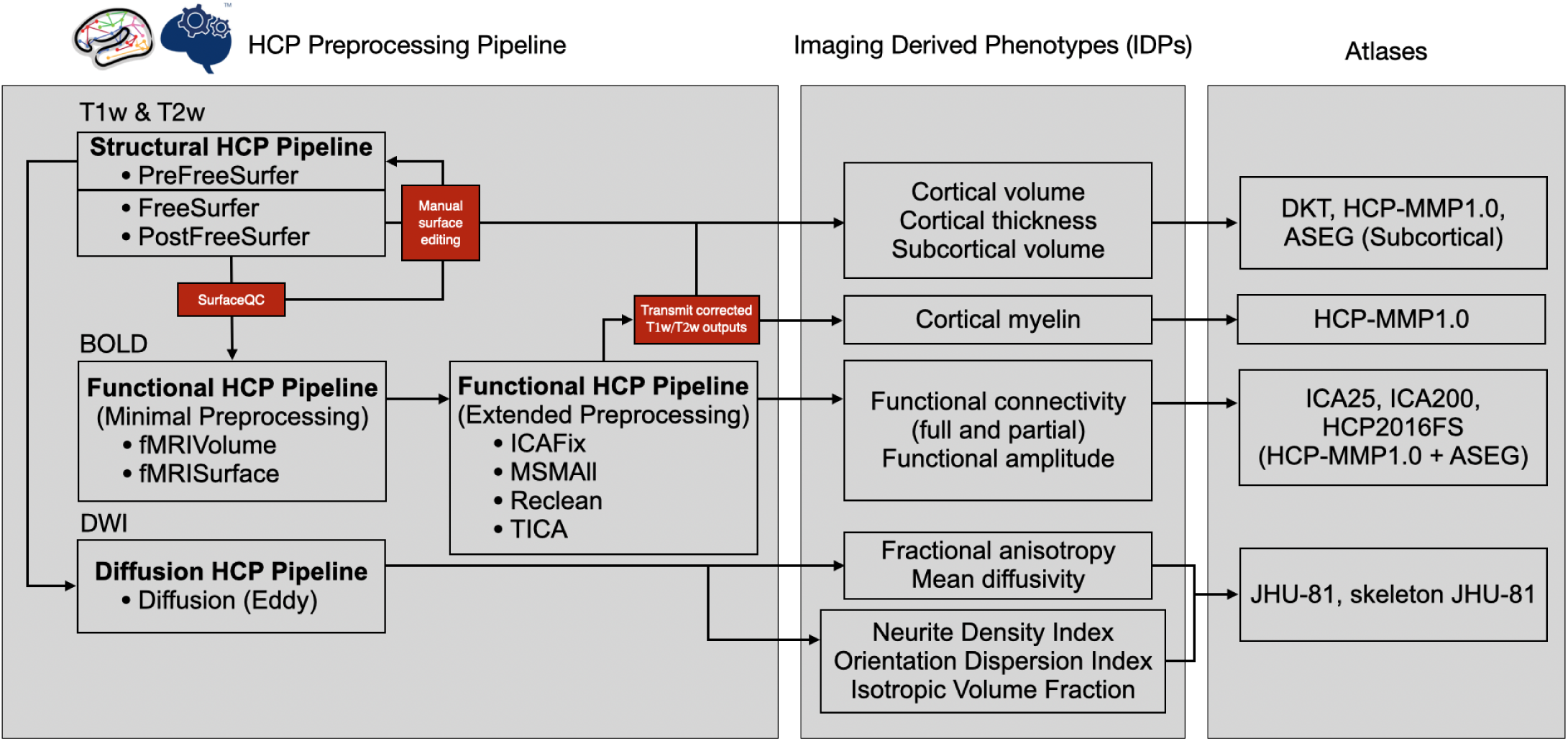
Flowchart overview of HARMONY processing pipeline. Names in the preprocessing pipeline on the left refer to modules in QuNex^72^. More information on IDP extraction and atlases can be found below. HCP = Human Connectome Project; QC = quality control; ICA = independent component analysis; MSM = multimodal surface matching; TICA = temporal ICA; DKT = Desikan-Killiany-Tourville; MMP = multimodal parcellation; ASEG = Automatic subcortical segmentation; JHU = Johns Hopkins University.

For quality control (QC), we evaluated signal-based and motion-related metrics, along with additional modality-specific measures, across all three imaging modalities. QC profiles across cohorts are shown in Fig. 3, with some expected variation in signal-based metrics. For example, HCP-DES showed higher signal-based metrics for sMRI and fMRI yet lower metrics for dMRI, which can be influenced by the difference in MRI hardware and acquisition protocols (see Supplementary Table S2). Case-control comparisons were performed for all QC measures separately per site. Although some QC differences between cases and controls were observed (Supplementary Table S4), these were not consistent across modalities or across sites. The total number of ICA components was higher in the HCP-BANDA adolescent dataset than in the three adult datasets (Supplementary Fig. S2) as is commonly observed in younger cohorts^73^, but the ratio of noise to total ICA components was stable across datasets such that data quality of post-processed data is not expected to differ from the adult cohorts. Although the extended ‘Reclean’ pipeline step had minimal impact on most datasets, it resulted in reclassification of some components in the HCP-DES dataset (Supplementary Fig. S2). Subject specific quality control information for all measures are included in the HARMONY data resource to enable users to set study-specific QC-based inclusion/exclusion thresholds.

**Figure 3:**
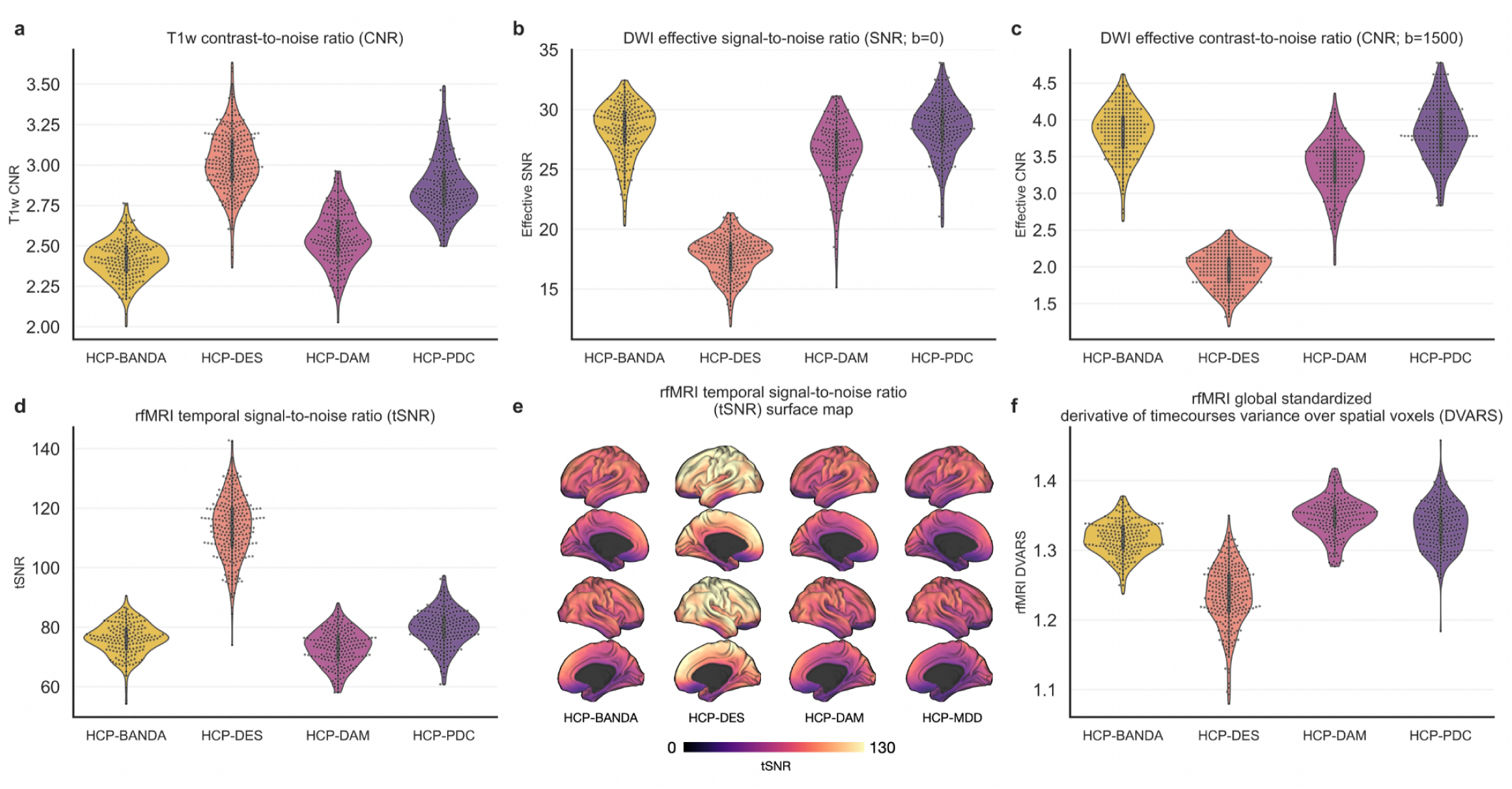
Quality control results on HARMONY data. For T1-weighted scans, the contrast-to-noise ratio (CNR) achieves scores above 2 for all HARMONY participants (a). For diffusion weighted imaging (DWI), effective signal-to-noise ratio (SNR) is reported for both b=0 (b) and b=1500 s/mm^2^ volumes (c). For rsfMRI, temporal signal-to-noise ratio (tSNR) is reported averaged across the grayordinates (d) and on the brain surface (e) for each CRHD cohort. Furthermore, DVARS (i.e., the root mean square of the temporal derivative) calculated from rsfMRI data and standardized with individual global signals is shown in the bottom right (f). Results in this figure are based on both cases and controls. Case-control comparisons for key QC measures can be found in Supplementary Table S4 and additional ICA-based QC measures for resting state fMRI and registration-based QC measures are shown in Supplementary Fig. S2.

### Extraction and harmonization of Imaging Derived Phenotypes (IDPs)

Following preprocessing and quality control, we extracted a total of 6 families of IDPs (2 families from each imaging modality; sMRI, rfMRI, and dMRI) with 13 unique IDP types across multiple atlases (Table 1). Briefly, IDP types from structural T1-weighted MRI included gray matter volumes, cortical thickness, and myelin estimated from the T1w/T2w ratio at the regional level using several different atlases (DKT & HCP-MMP1.0 for cortical areas and ASEG for subcortical regions^74,75^). IDP types from dMRI included both image-level and tract-level for fractional anisotropy, mean diffusivity, and neurite-related white matter microstructures. For rsfMRI, we included ROI- and network-based parcellations, covering both low and high feature dimensions, to enable systematic evaluation at the level of aggregated networks and cortical regions. Specifically, IDP types from rsfMRI included amplitudes (temporal standard deviation), full correlation connectivity, and partial correlation connectivity for two different atlases (ICA-25 covering large-scale networks and HCP-MMP1.0 covering 360 cortical areas combined with 19 subcortical regions from the ASEG atlas). Partial correlation connectivity was also calculated after combining bilateral homologues from the HCP-MMP-1.0 atlas to avoid overcorrection for bilateral synchrony. In summary, these IDP types comprise region-level tabulated summary measures and were selected to align with IDP families available in other large-scale datasets^76^. For each of the 13 IDP types covering all three imaging modalities, we selected a representative atlas as a focus for all subsequent analysis. The selected atlases for each IDP type are highlighted with (^d^) in Table 1.

**Table 1:** HARMONY Imaging Derived Phenotypes (IDP)

Imaging derived phenotypes (IDPs) capture tabulated summary information regarding brain structure and function for parcellated brain regions.
| IDP family | IDP type | Atlas | IDP measures | Number of used features |
| --- | --- | --- | --- | --- |
| Brain Morphometry | Gray matter volume | Desikan-Killiany-Tourville (DKT) | cortical_volume_DKT | 63 |
|  |  | HCP-MMP1.0 (Glasser) | cortical_volume_HCP-MMP | 360 |
|  |  | HCP-MMP1.0 and 19 Freesurfer Subcortical Segmentation ASEG (HCP2016FS) | volume_HCP2016FS<sup>d</sup> | 379<sup>b</sup> |
|  |  | Freesurfer Subcortical Segmentation (ASEG, 19 regions) | subcortical_volume_FS | 19<sup>a</sup> |
|  | Cortical thickness | Desikan-Killiany-Tourville (DKT) | cortical_thickness_DKT | 63 |
|  |  | HCP-MMP1.0 (Glasser) | cortical_thickness_HCP-MMP<sup>d</sup> | 360 |
| Myelin | Cortical myelin | HCP-MMP1.0 (Glasser) | cortical_myelin_HCP-MMP<sup>d</sup> | 360 |
| Functional Amplitude | Functional amplitude | HCP-MMP1.0 and 19 ASEG (HCP2016FS) | amp_HCP2016FS<sup>d</sup> | 379 |
|  |  | HCP-YA-derived 25 Independent Component Analysis (ICA25) | amp_ICA25 | 25 |
|  |  | HCP-YA-derived 200 Independent Component Analysis (ICA200) | amp_ICA200 | 200 |
| Functional Connectivity | Full connectivity | HCP-MMP1.0 and 19 ASEG (HCP2016FS) | full_corr_HCP2016FS | 71,631<sup>c</sup> |
|  |  | HCP-YA-derived 25 Independent Component Analysis (ICA25) | full_corr_ICA25<sup>d</sup> | 300<sup>c</sup> |
|  |  | HCP-YA-derived 200 Independent Component Analysis (ICA200) | full_corr_ICA200 | 19,900<sup>c</sup> |
|  | Bilateralized full connectivity | HCP-MMP1.0 and 19 ASEG (HCP2016FS) | bilat_full_corr_HCP2016FS<sup>d</sup> | 17,955<sup>c</sup> |
|  | Partial connectivity | HCP-MMP1.0 and 19 ASEG (HCP2016FS) | partial_corr_HCP2016FS | 71,631<sup>c</sup> |
|  |  | HCP-YA-derived 25 Independent Component Analysis (ICA25) | partial_corr_ICA25<sup>d</sup> | 300<sup>c</sup> |
|  |  | HCP-YA-derived 200 Independent Component Analysis (ICA200) | partial_corr_ICA200 | 19,900<sup>c</sup> |
|  | Bilateralized partial connectivity | HCP-MMP1.0 and 19 ASEG (HCP2016FS) | bilat_partial_corr_HCP2016FS<sup>d</sup> | 17,955<sup>c</sup> |
| Diffusion-tensor white matter microstructure | Fractional anisotropy (FA) | Johns Hopkins University White-Matter Labeled (JHU-81) | fa_JHU81 | 48 |
|  |  | HARMONY-derived skeleton with Johns Hopkins University White-Matter Labeled (JHU-81) | skeleton_fa_JHU81<sup>d</sup> | 48 |
|  | Mean diffusivity (MD) | Johns Hopkins University White-Matter Labeled (JHU-81) | md_JHU81 | 48 |
|  |  | HARMONY-derived skeleton with Johns Hopkins University White-Matter Labeled (JHU-81) | skeleton_md_JHU81<sup>d</sup> | 48 |
| Neurite-related white matter microstructure | Neurite density index (NDI) | Johns Hopkins University White-Matter Labeled (JHU-81) | ndi_JHU81 | 48 |
|  |  | HARMONY-derived skeleton with Johns Hopkins University White-Matter Labeled (JHU-81) | skeleton_ndi_JHU81<sup>d</sup> | 48 |
|  | Orientation dispersion index (ODI) | Johns Hopkins University White-Matter Labeled (JHU-81) | odi_JHU81 | 48 |
|  |  | HARMONY-derived skeleton with Johns Hopkins University White-Matter Labeled (JHU-81) | skeleton_odi_JHU81<sup>d</sup> | 48 |
|  | Isotropic volume fraction (IVF) | Johns Hopkins University White-Matter Labeled (JHU-81) | ivf_JHU81 | 48 |
|  |  | HARMONY-derived skeleton with Johns Hopkins University White-Matter Labeled (JHU-81) | skeleton_ivf_JHU81<sup>d</sup> | 48 |

Although all image acquisition sequences were inspired by HCP-style protocols, site effects may arise from scanner model, acquisition-parameter differences, and other site-specific factors, even when protocols are closely matched (Supplementary Table S2). To account for these effects, we conducted IDP harmonization using ComBat-GAM^63^ and CovBat-GAM^77^. The current implementation of ComBat-GAM allows for training on controls only and applying the trained adjustments to cases^78^, which is important given that symptom severity systematically differed between sites. CovBat-GAM does not currently have this functionality, and we therefore included a measure of anhedonia as a covariate in the implementation of CovBat-GAM in order to retain variance related to affective symptoms during the CovBat-GAM harmonization. The impact of IDP harmonization was assessed using site classification accuracy based on multiclass support vector machines and illustrated by association curves between gray matter volume and age. Notably, low accuracy in site classification is desirable as it indicates successful harmonization.

Qualitatively, the associations between age and overall gray matter volume showed undesired offsets between cohorts prior to harmonization (Fig. 4a) that were effectively removed by applying CovBat-GAM (Fig. 4b), and ComBat-GAM (Fig 4c). As expected, site classification was significantly higher than chance in both cases and controls for all unharmonized IDPs (Fig. 4d-e; Supplementary Fig. S3). After applying CovBat-GAM and ComBat-GAM, site-related effects in the IDPs were attenuated in the control group, as reflected by a marked reduction in site-prediction accuracy (Unharmonized: 77%; CovBat-GAM: 38%; ComBat-GAM (fit on controls): 31%; Fig. 4). In contrast, the attenuation of site classification accuracy for cases showed greater differences between ComBat-GAM (fit on controls) and CovBat-GAM. Specifically, in cases ComBat-GAM resulted in a site classification accuracy of 71%, whereas accuracy in CovBat-GAM was lower at 38% (compared to 83% prior to harmonization). Importantly, the intention of harmonization is to remove site effects while retaining meaningful individual differences. As such, the ComBat-GAM pattern of low site classification accuracy in controls yet high site classification accuracy in cases suggests that, in the case group, site-related differences are partially confounded with clinical differences across cohorts. Although we included severity as a covariate to preserve in CovBat-GAM, our findings demonstrate that the inclusion of patients in the model may nevertheless result in over-correction. On the other hand, training ComBat-GAM on controls and applying the estimated adjustments to cases may preserve clinically relevant signals, and therefore does not minimize site classification accuracy among cases due to overlap with a possible anhedonia effect. The effects of ComBat-GAM versus CovBat-GAM were assessed further in the context of brain-symptom associations as described below.

**Figure 4:**
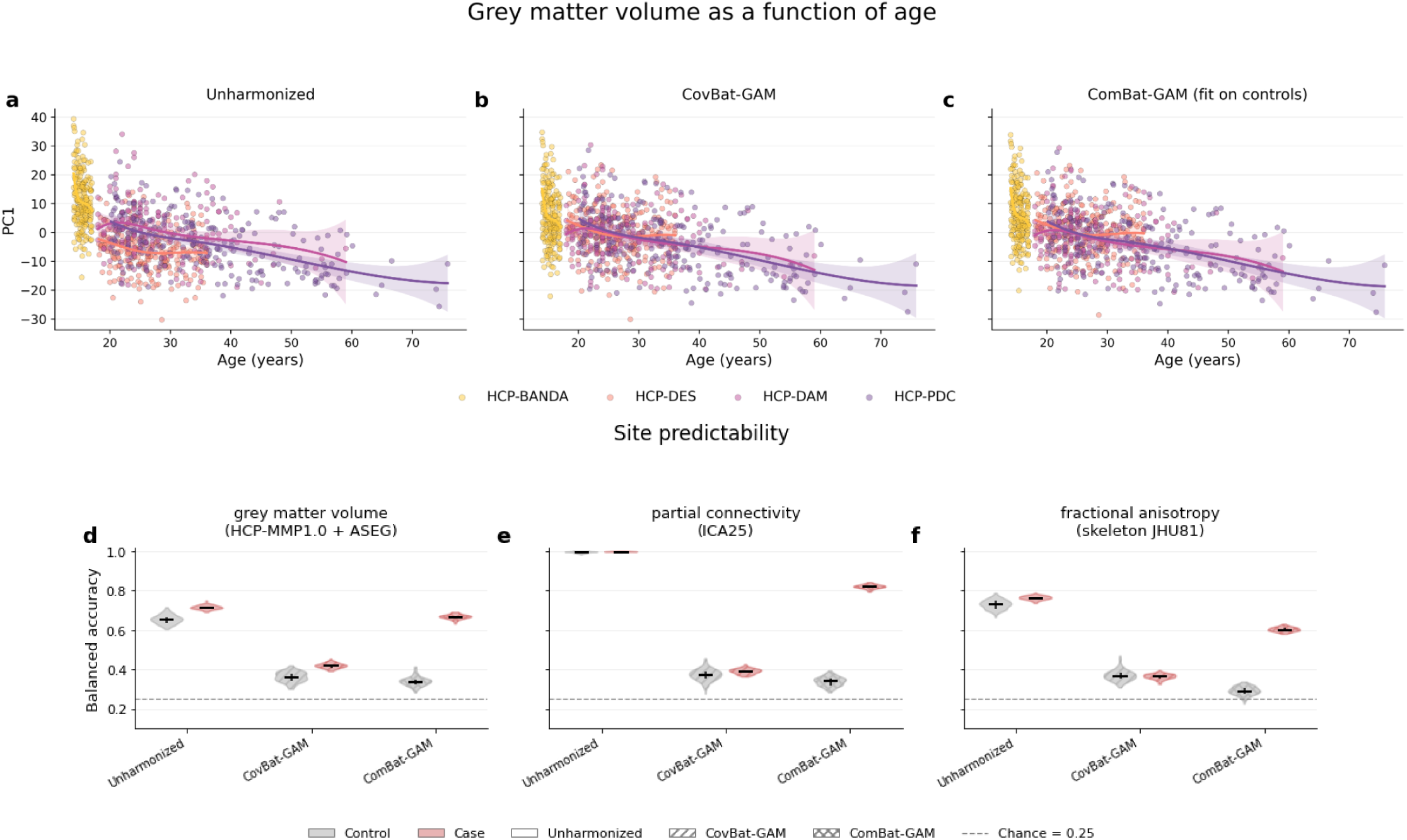
Harmonization strategies were illustrated by the relationship with principal component 1 (PC1) of gray matter volume as a function of age for unharmonized (a), CovBat-GAM (b), and ComBat-GAM (c). Specifically, only cases were illustrated and age was modeled using a B-spline basis with 5 degrees of freedom and cubic spline, and PC1 was regressed on this spline basis using ordinary least squares. 95% confidence intervals were obtained from the model for each cohort: HCP-BANDA (yellow), HCP-DES (orange), HCP-DAM (ruby), and HCP-PDC (purple). Moreover, IDP site effects were evaluated by cross-validated SVM site classification before and after harmonization (lower accuracies reflect greater site-correction). Balanced accuracy was estimated using PCA (k=25) with 100 repeated 5-fold cross-validation. Results showed effective harmonization (i.e., removal of site classification accuracy) for both ComBat-GAM (fit on controls) and CovBat-GAM in controls (gray) for three selected IDPs: gray matter volume (d) partial connectivity ICA25 (e), and skeletonized fractional anisotropy (f). Elevated site classification accuracy in ComBat-GAM (fit on controls) for cases only (red) was presumably driven by systematic differences in psychopathology across sites, reflecting retention of true site differences that are of interest. The dashed line indicates chance-level balanced accuracy. Here, and in the other figures, the black vertical bars in the distribution plots indicate the interquartile range and the black horizontal bars indicate the median.

### Brain-symptom associations

To provide a proof-of-concept analysis of brain–behavior associations in the HARMONY dataset, we quantified associations between 13 selected IDPs and (i) anhedonia (measured using the Snaith-Hamilton pleasure scale (SHAPS)^79^) and (ii) depression severity (measured using the Hamilton depression rating scale (HAMD)^80^). Anhedonia analyses included all four cohorts, whereas depression severity analyses excluded HCP-BANDA because HAMD was not acquired in this cohort. Only cases were included for subsequent analyses. Multivariate prediction analyses involved principal component regression (PCR) with 100 repeats of 5-fold cross-validation (with PCA and covariate regression performed within folds to avoid leakage). Here, PCR was chosen to match the input dimensionality across all IDPs, and the size of the reduced feature space was set to k=25 to match our smallest IDP feature set. Univariate analyses involved separate partial correlation per feature, adjusted by age and sex, followed by false discovery rate correction for multiple comparisons within each IDP.

Across 13 selected IDP measures, ComBat-GAM achieved significant cross-validated R^2^ for the prediction of anhedonia scores when using structural and functional IDPs, but not for diffusion IDPs (Fig. 5 and Supplementary Fig. S4). When comparing multivariate predictive performance as a function of harmonization approaches, ComBat-GAM (fit on controls) consistently outperformed CovBat-GAM (Fig. 5). These results suggest that relatively more variance relevant to affective symptoms was retained when training on healthy controls only and applying to cases. This feature is currently only implemented in ComBat-GAM, and we anticipate improved results for CovBat-GAM harmonization once the same approach can be applied. Accordingly, we used ComBat-GAM–harmonized IDPs for subsequent univariate analyses. Prediction accuracy also varied across IDP types (Fig. 5b).

**Figure 5:**
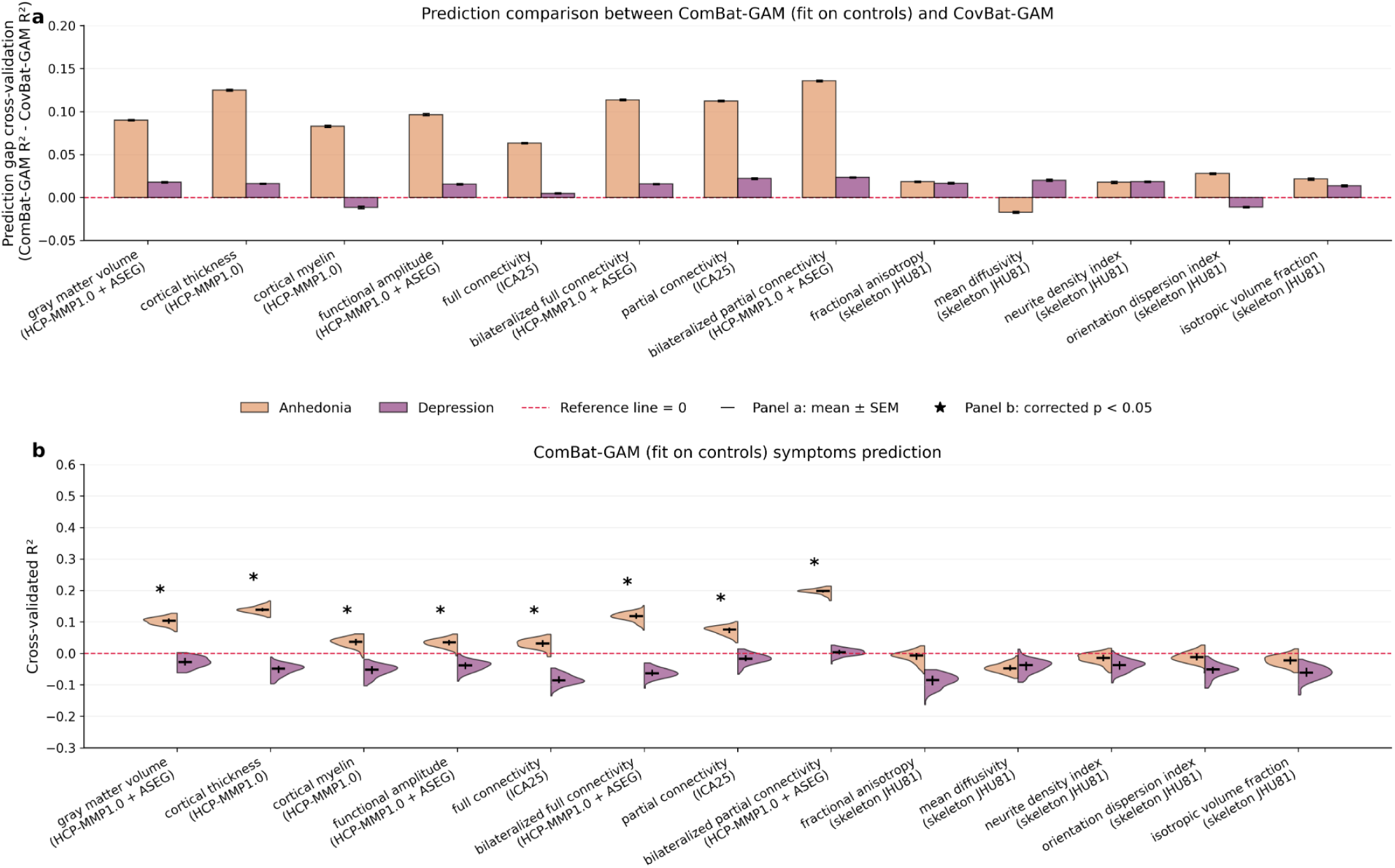
Multivariate brain-symptom association results. Principal component regression (k=25) with 100 repeated 5-fold cross-validation were performed after adjusting for age and sex at feature level. (a) Out of the 13 IDP types (with one pre-selected atlas per type), R^2^ was higher for ComBat-GAM (fit on controls) than for CovBat-GAM in almost all analyses (12 out of 13 for anhedonia, and 11 out of 13 for depression severity). Bars show the difference in R2, with positive values indicating higher R^2^ for ComBat-GAM (fit on controls), and error bars show the standard error of mean. (b) Out of 13 IDP types (with one pre-selected atlas per type), bilateralized functional connectivity achieved the highest R^2^ for anhedonia (orange) but showed no significant effect for depression severity (purple) prediction. Distributions in (b) represent the variation across 100 repetitions of 5-fold cross-validation (with R^2^ averaged across folds within a repetition). Note that R^2^ is calculated from the sum squared error (rather than the squared correlation) and can therefore take on negative values indicative of poor model performance. The asterisks refer to significant predictions (p < 0.05) calculated from a corrected resampled Student’s t-test against zero^118^. Anhedonia results (orange) included all 4 cohorts and depression results (purple) included only the 3 adult cohorts because HAMD was not available in HCP-BANDA. Results in this figure are based on cases only.

To assess the value of the combined HARMONY dataset compared to individual cohorts, we compared model performance when training and testing models in each separate dataset to the performance when training and testing models on the aggregated HARMONY dataset. As performance metrics, we used overall prediction accuracy and the gap between training and testing R^2^ values (i.e., an indicator of the improved cross-validated generalizability reflected by reduced overfitting of estimates). Cross-validated R^2^ prediction accuracy for anhedonia reached significance for the HARMONY dataset – but not for individual cohorts – when trained on gray matter volume and functional partial connectivity IDPs (Fig. 6a-b), but not diffusion IDP (fractional anisotropy; Fig. 6c). In addition to significant cross-validated prediction performance, the gap between training and test accuracy was substantially lower in the HARMONY dataset compared to individual cohorts (Fig. 6d), indicating improved performance and cross-validated generalizability in the combined dataset. Repeating the analyses in Fig. 6 after mixing subjects across cohorts revealed that both the prediction accuracy and train-test gap advantage of HARMONY was explained by increased sample size, rather than cohort mixing (Supplementary Fig. S5).

**Figure 6:**
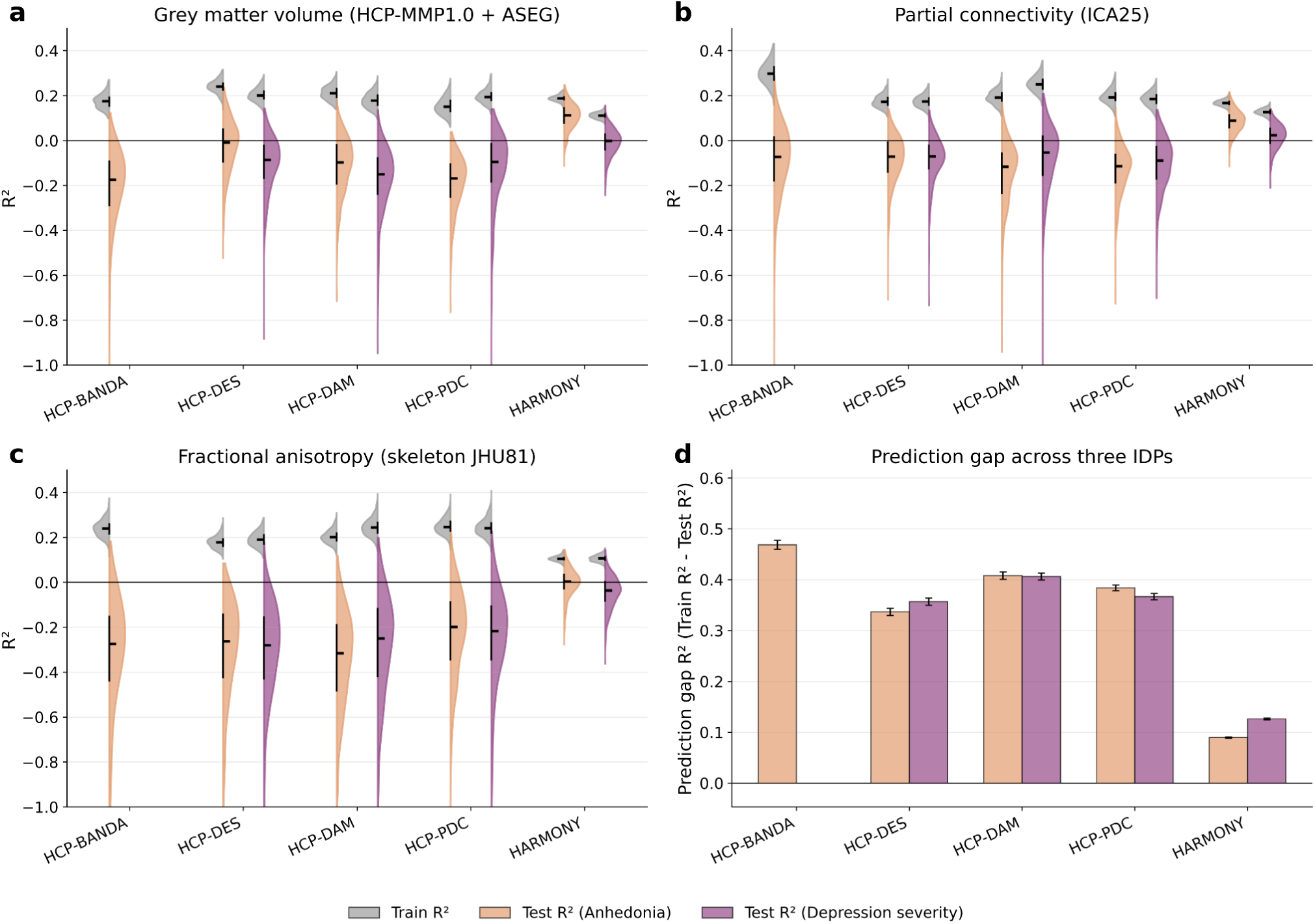
Comparison of multivariate brain–symptom models trained either on separate CRHD cohorts or on the combined HARMONY dataset for three ComBat-GAM-harmonized IDPs, including gray matter volume (a), partial connectivity (b) and fractional anisotropy (c). Prediction gaps (train R^2^ - test R^2^) were calculated for each of 100 repetitions, and averaged across the three selected IDPs shown in a, b, and c (d). Error bars indicate standard errors of mean across 300 datapoints (3 IDPs x 100 repetitions each). Results indicated reducing overfitting to testing data, reflected by smaller prediction gaps between train and test splits for the combined dataset relative to the individual cohorts. Anhedonia results (orange) were available for all 4 cohorts and depression severity results (purple) were not available in HCP-BANDA. Results in this figure are based on cases only.

For univariate results, we focused on associations of anhedonia and depression severity, with three ComBat-GAM-harmonized IDPs representative of structural, functional, and diffusion modalities. Out of 379 gray matter volume IDP features, 36 IDP features passed false discovery rate (FDR) correction in relation to anhedonia (Fig. 7a). Specifically, significant gray matter volume reductions associated with anhedonia were observed in both left and right hemispheres spanning frontal, middle temporal, and parietal regions. Notably, cingulate involvement was also present in classic emotion-related circuits, including anterior/mid cingulate and posterior cingulate/retrosplenial regions. Within significant features, two left cortical and one subcortical regions showed positive associations (including left lateral occipital, left parieto-occipital sulcus, and right ventral diencephalon). Effect sizes were generally modest, with an average correlation of r = -0.12 among negative associations. On the other hand, none of the depression severity associations passed FDR correction after controlling for age and sex. The regions with p < 0.05 uncorrected are shown in gray in Fig. 7b.

**Figure 7:**
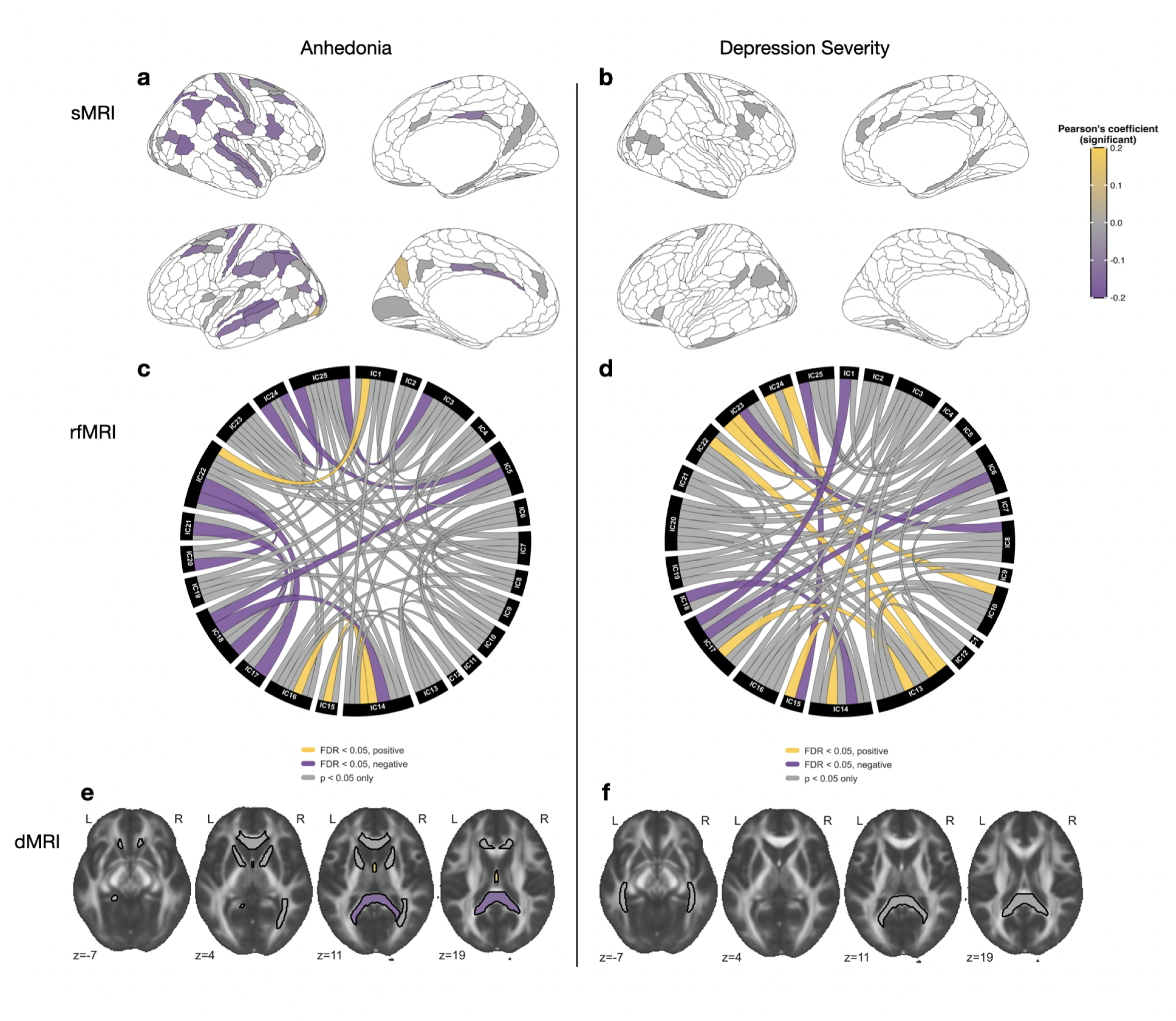
Univariate associations between imaging phenotypes and anhedonia (left column) or depression severity (right column). Effect sizes of gray matter volume with anhedonia (a) and depression severity (b) were estimated by partial correlation (regressed out age and sex) with Benjamini-Hochberg false discovery rate (FDR). Regions with p_fdr_ < 0.05 are plotted in color either purple (negative association) or yellow (positive association), while regions with p < 0.05 uncorrected are shown in gray. Chord diagrams were visualized for association between partial functional connectivity (ICA25) with anhedonia (c) and depression severity (d). FMRIB58 was used as a template to project relationships between fractional anisotropy (skeleton JHU81) and depression (e) or anhedonia (f). Results in this figure are based on cases only.

For rsfMRI, the low-dimensional representation of 25-dimensional ICA enabled interpretability of partial functional connectivity. The results showed both positive and negative associations with anhedonia (Fig. 7c) with the average absolute magnitude around 0.15 across 11 significant FDR-corrected edges. Using the 25-component sICA decomposition (see Supplementary Fig. S6a for template ICA maps), negative associations with anhedonia were primarily observed among connections between default-mode network, fronto-parietal network, temporo-parietal junction, visual network, thalamus, precuneus, caudate nucleus and cerebellum while positive association with anhedonia were observed in among visual network, temporo-parietal junction and cerebellar connections. On the other hand, depression severity showed average absolute magnitude around 0.14 across 11 significant FDR-corrected edges (Fig. 7d). Six positive associations were observed among fronto-parietal network, tempero-parietal junction, somatosensory network, visual network, caudate nucleus and cerebellar connections, while five negative associations primarily observed among default-mode network, fronto-parietal network, tempero-parietal junction, visual network, precuneus, cerebellum and brain stem connections.

For dMRI, higher anhedonia was associated with lower fractional anisotropy in the splenium of the corpus callosum (r=-0.12, p_fdr_ = 0.027) and higher fractional anisotropy in the fornix (r=0.11, p_fdr_ = 0.027). Depression severity showed no significant dMRI associations after correction (see Supplementary Fig. S6b for template skeleton map).

### Data sharing

The HARMONY data resource will shortly be available for download at the NDA and at BALSA. The data resource includes easily accessible tabulated data covering multimodal IDPs (Table 1), symptom data, demographics, quantitative QC metrics, and a data dictionary.

Data access steps are akin to HCP-YA access and require all users to agree to Data Use Terms. Although the HARMONY data is deidentified, combinations of variables could be potentially reidentifying. As such, it is important that users consult with an Institutional Review Board or Ethics Committee before beginning the research to ensure that appropriate approvals/exemptions are in place.

Tabulated data are shared for all participants for whom processing pipelines reached completion. Future studies using the data resource may wish to leverage QC metrics to exclude participants with noisy data, while assessing and balancing the risk of bias^81^. To this end, we have shared tabulated subject-level information on all QC measures described above. We elected to share ‘raw’ (unharmonized) IDPs and ComBat-GAM (fit on controls) harmonized IDPs. The corresponding scripts were also provided, allowing users to rerun the harmonization pipeline and modify the hyperparameters as needed for CovBat-GAM harmonization and their own specific analyses.

## Discussion

HARMONY was developed to support robust and reproducible neuroimaging research on anxious misery by integrating multiple state-of-the-art HCP-style cohorts into a single, harmonized, multimodal resource. Large-scale harmonized neuroimaging initiatives have demonstrated transformative impact across the full research landscape from standardizing acquisition and preprocessing^85^, to enabling disease-focused discovery and benchmarking^86–88^ and linking brain measures to genetics^89–91^. Large scale datasets also provide the sample sizes required for modern data-driven approaches, including machine learning and deep learning, where model robustness improves with larger, diverse, and well-curated data^92,93^. In this context, HARMONY extends these big data advantages to anxious misery research.

Importantly, our findings showed that pooling harmonized datasets improved statistical power for detecting subtle effects in the neurobiological correlates of anxious misery. Effect sizes were larger in the combined HARMONY dataset compared to the individual cohorts, with improved generalizability to unseen data and improved statistical power. Importantly, HARMONY offers a clinically curated and diagnostically stratified sample that has been phenotyped using robust clinician-rated instruments (e.g., HAMD), offering rich opportunities for research into heterogeneity and subtypes of anxious misery^11^.

When combining multiple datasets, a key challenge is to implement appropriate control for site differences, while retaining effects of interest. Our findings showed that both ComBat-GAM (fit on controls) and CovBat-GAM successfully reduced site effects across all released IDPs. Importantly, the severity of mental health symptomatology varied systematically across HARMONY sites, which may lead to the attenuation of severity effects after harmonization if not carefully handled^94,95^. To avoid unintentional removal of severity effects, we compared (i) the CovBat-GAM approach^62^ used in a prior data resource^77^ and implemented to retain variance linked to covariates of interest (here age, sex, and SHAPS)^63^ with (ii) an approach, ComBat-GAM, that can be trained on just healthy controls, with the estimated site adjustments then applied to the “unseen” case data. Our findings revealed stronger associations between IDPs and anhedonia after the ComBat-GAM harmonization (fit on controls), with appropriately attenuated site effects in controls. Notably, site effects were not fully removed among cases following ComBat-GAM, likely reflecting true site-related differences in the severity of anxious misery symptoms. These results reveal the tension between desirable site correction and undesirable removal of effects of interest. Notably, functionality to train CovBat-GAM on controls and apply to unseen case data is underway at the time of writing. Once available, this and other harmonization techniques that enable control-only fitting may be suitable for further exploration in the HARMONY data resource.

In terms of brain-behavior associations in the anxious misery domain, our findings demonstrated that functional IDPs achieved the strongest multivariate predictive performance across the three imaging modalities and across IDP families. Recent benchmarking work suggests that properties of resting-state connectivity maps depend strongly on the chosen connectivity metric, with partial connectivity achieving stronger individual fingerprint predictability and higher mental health predictability compared with standard full connectivity^96^. Furthermore, partial connectivity achieves improved correspondence between structural and functional connectivity as compared to full connectivity^97^. As such, partial functional connectivity IDPs offer a particularly promising representation for future transdiagnostic prediction and biomarker-oriented analyses.

In addition to its strengths, including a harmonized sample of 743 individuals with anxious misery spanning a broad age range and symptom severity range, several limitations should be considered when interpreting the HARMONY resource. First, although all four CRHD cohorts acquired extensive batteries to characterize anxious misery symptomatology^28^ only HAMD and SHAPS were consistently available across (adult) cohorts. This constrains the range and specificity of harmonized symptom domains, which are necessarily reduced to constructs that can be aligned across datasets. Future work may extend this by integrating partially overlapping instruments through data-driven or latent variable approaches. Second, although quality-control (QC) profiles are provided, variability in data quality remains an important consideration, particularly given its influence on brain-behavior associations^98,99^. Further work could incorporate more advanced QC metrics and systematically evaluate the impact of subject-level exclusion thresholds on downstream analyses^81,100^, as well as the role of natural quality variation relevant to future clinical translation in real-world settings that use standardized QC routines. Third, the interpretation of brain–symptom associations in large-scale datasets depends critically on the nature of the sampled variation. Analyses that span broad ranges of symptom severity, including subclinical levels, may capture low-dimensional or global patterns of variation but may also dilute associations that are more specific to stratified subgroups or higher severity presentations^15,47,101,102^. Although HARMONY represents an advance over general population datasets through its focus on samples specifically recruited for active symptomatology, the distribution of severity remains predominantly in the mild-to-moderate range. These considerations highlight both the strengths and the appropriate scope of the HARMONY resource: it is well suited to population-level, cross-cohort analyses of broad brain–behavior associations, while complementary approaches that more directly parse heterogeneity—through targeted phenotyping, subgroup identification, or specific circuit-informed measures—will be important for resolving more specific and clinically relevant patterns of brain dysfunction.

In summary, we developed the HARMONY resource by harmonizing four CRHD datasets through standardized HCP-style preprocessing, rigorous quality control, and the systematic extraction and integration of imaging derived phenotypes and symptom measures. HARMONY is designed to support reproducible, cross-cohort investigation of brain-behavior relationships in anxious misery, particularly through data-driven analyses of population-level variation. We anticipate that this resource will facilitate studies of heterogeneity, subgroup stratification, dimensional symptom variation, and lifespan effects, providing a foundation for more targeted investigations of brain circuit function in depression and anxiety.

## Methods

### Participants

Secondary data were aggregated from four CRHD cohorts that acquired multimodal HCP-style protocols and shared de-identified imaging, clinical and behavioral data^28^. HCP-BANDA included 152 adolescents aged 14–17 years, including youth with a diagnosis of depression and/or anxiety. HCP-DES included 222 symptomatic, untreated young adults aged 18–36 years; clinical inclusion required clinically significant anhedonia, anxious arousal, concentration difficulties, rumination, or tension. HCP-DAM included 197 participants with anxious misery symptoms, with an inclusion criterion of neuroticism ≥1 standard deviation above the mean of controls. Finally, HCP-PDC comprised 172 adults aged 20–76 years with major depressive disorder who failed to respond to at least two antidepressants, with no neuromodulation or ketamine within the previous 6 months. All sites included healthy controls (N=49-63 additional per site) who did not meet DSM-5 criteria for psychiatric disorders (See Supplementary Table S1 for a detailed breakdown of cohort characteristics).

### Longitudinal designs

While this study focuses mainly on baseline characteristics with available neuroimaging data, all four CRHD included a follow-up component, which involved non-imaging measurements for HCP-BANDA,HCP-DES, and HCP-DAM, and a full protocol repeat following treatment interventions for HCP-PDC. More information on longitudinal designs is provided in the supplementary information. IDPs derived from follow-up imaging sessions are included in the data release.

### Collation of demographic and symptom data

To collate non-imaging data across HARMONY sites, we first isolated and formatted each instrument into a stand alone comma-separated values (CSV) file mapped to an NIMH Data Archive (NDA) “structure” wherever possible (typically but not always 1:1 at the instrument level). These files served as the basis for a harmonized cross-site data dictionary, which we developed in several stages. First, we identified instruments with differing or missing names and descriptions (e.g., `hrds01` and `hamd01` both representing the HAMD^80^), and linked instruments using a common shortname (`hamd01`) and instrument label. Next, we isolated and harmonized key demographic variables (age, sex, race, ethnicity, adapted GUID, collection date) using NDA dictionary definitions (e.g., age recorded in months rather than years). We then removed housekeeping or redundant variables (e.g., `form_complete` fields and duplicate subject identifiers) and merged MR session identifiers.

We additionally harmonized identically collected variables with different names by creating and storing explicit rename maps. Because behavioral assessments were often collected across multiple adjacent days, the `interview_date` could differ across components, producing multiple sparse records per participant. We distinguished scientifically meaningful longitudinal spacing (e.g., quarterly visits and pre-post treatment visits) from practical scheduling windows (e.g., 2–3 adjacent days) and collapsed multi-day records into a single visit aligned to the imaging session where appropriate, using the earliest date as the visit date.

We then conducted targeted checks for common issues encountered when integrating behavioral and clinical measures across cohorts, including (i) inconsistent scale coding (e.g., Likert scales coded 0–4 vs. 1–5), (ii) reverse coding (0–4 vs. 4–0), (iii) differences in summary-score computation, (iv) differences in missing-value encodings, and (v) distributional shifts indicative of procedural or platform changes or cohort differences. The resulting shared HARMONY data resource includes core demographic variables (age, sex, race, ethnicity, collection data) and symptom data leveraged in this study (case/ control status, HAMD, and Snaith-Hamilton Anhedonia Scale (SHAPS)^79^).

SHAPS and HAMD had different levels of missingness across cases and controls. Among cases (n = 743), SHAPS was available for 698 participants and missing for 45 participants (6.1% missing), while HAMD was available for 523 participants and missing for 68 participants (11.56% missing from 3 available datasets). Among controls (n = 220), SHAPS was available for 207 participants and missing for 13 participants (5.9% missing), while HAMD was available for 131 participants and missing for 26 participants (16.56% missing). At the site level, SHAPS missingness among cases was highest in HCP-DAM (37/197; 18.8%), followed by HCP-BANDA (4/152; 2.6%) and HCP-PDC (4/172; 2.3%), with no missing SHAPS data in HCP-DES cases. For HAMD, missingness among cases was highest in HCP-DAM (39/197; 19.8%), followed by HCP-DES (27/222; 12.2%) and HCP-PDC (2/172; 1.2%).

### MRI acquisition

MRI acquisitions were designed to follow the HCP-style protocols, with three studies acquired on 3T Siemens Prisma systems (HCP-BANDA, HCP-DAM, HCP-PDC) and one on 3T GE Discovery MR750 or UHP (HCP-DES), using high-density receive-array head coils and comparable sequence timing overall. Full acquisition parameters are available in Supplementary Table S2, and cross-site comparisons are described in detail by Tozzi et al^28^. In brief, all studies collected high-resolution 0.8 mm isotropic 3D T1w and T2w scans with similar field-of-view and contrast settings. For both fMRI and dMRI, the EPI acquisitions included runs with both “AP” and “PA” phase encoding polarity, with AP/PA pairs of spin-echo EPI images also acquired to support susceptibility distortion correction. Resting-state fMRI used highly accelerated multiband EPI (TR = 0.8 s for Siemens; 0.71 s for GE), but HCP-DES differed by using MB=6 (vs MB=8) and 2.4 mm voxels (vs 2.0 mm). Diffusion MRI was similarly harmonized across sites using multi-shell b=1500/3000 s/mm² protocols (MB=4 with 1.5 mm isotropic voxels), although HCP-DES acquired fewer directions per shell (75/75 vs 92/93 for the Siemens cohorts). Total duration of the resting-state scans varied by project, with HCP-DES/DAM/BANDA acquiring ∼20 minutes per session, whereas HCP-PDC acquired ∼12 minutes per session.

### Image preprocessing

All datasets were processed using the HCP Pipelines as implemented in the QuNex container (see Supplementary Table S5)^24,64,65^. For structural and functional MRI, the minimal preprocessing workflow included PreFreeSurfer, FreeSurfer, and PostFreeSurfer for anatomical reconstruction and surface generation, followed by fMRIVolume and fMRISurface for resting-state fMRI preprocessing projected to CIFTI grayordinates space. Surface-based registration quality was assessed by manual review of cortical surface placement and “myelin map” quality, and scans were flagged for reprocessing when clear surface or registration imperfections were qualitatively observed (Supplementary Table S3). Cortical myelin estimates were derived from the transmit field–corrected T1w/T2w maps to mitigate spatial bias from B1+ transmit inhomogeneity^103^.

For resting-state fMRI, an extended denoising pipeline was applied to improve artifact removal beyond minimal preprocessing. This included spatial ICA with FIX for automated component classification and removal of structured noise^67^. We additionally applied ‘Reclean’ (recleaning) to refine ICA-based classification by re-evaluating components assignment and improving noise identification^70^. Finally, ‘Tclean’ (temporal ICA; tICA) was used to remove global and temporally structured artifacts while preserving neural signals^71,104^. Surface alignment across subjects was performed using multimodal areal-feature-based registration (‘MSMAll’) to improve correspondence of cortical areas^69^.

For diffusion MRI, data were processed using the HCP diffusion preprocessing workflow, including susceptibility-induced distortion correction using TOPUP and eddy-current correction using EDDY^105^, with rotation of diffusion gradient directions, outlier replacement (with `--ol_nstd`=5)^106^, movement-by-susceptibility correction^107^, and slice-to-volume (S2V) movement correction^108^. S2V was not included for HCP-DES, since the necessary slice-timing information of the General Electric multiband implementation was not available.

### Quality control measures

In addition to manual QC, we quantified signal-based and motion-related metrics, together with modality-specific QC measures, to provide a holistic view of neuroimaging data quality. Signal-based QC included contrast-to-noise ratio (CNR) for T1w (computed with Freesurfer), temporal signal-to-noise ratio (tSNR) for blood-oxygenation-level–dependent (BOLD) echo planar imaging (EPI), which was computed using FSL tools (e.g., fslmaths and fslstats), and SNR/CNR reported by QUAD for diffusion weighted imaging (DWI)^109^. Specifically, FreeSurfer’s ‘mri_cnr’ was used to estimate structural CNR by sampling intensities around the cortical surface (white matter, cortical gray matter, and CSF) and computing an averaged tissue contrast-to-noise measure (primarily gray–white and gray–CSF contrast terms) reflecting local tissue-intensity variability rather than image background.

For resting-state fMRI, tSNR was computed at each grayordinate/voxel as:

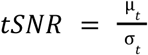

where μ_t_ is the mean signal across time and σ_t_ is the standard deviation across time; the resulting tSNR map was then averaged across grayordinates/voxels to obtain subject-level summary values. In addition to the signal-based metrics, we quantified the root mean square of the temporal derivative (DVARS) as a motion-related QC metric, reflecting frame-to-frame changes in whole-brain BOLD signal intensity that are often associated with head motion and other transient non-neural artifacts^110^.

For DWI, we used QUAD-reported SNR/CNR summary metrics from the FSL ‘eddy’ QC pipeline, where an SNR map was computed from the b=0 volumes, and CNR maps were calculated for each of the diffusion-weighted shells (b= 1500 and 3000 s/mm^2^) as follows:

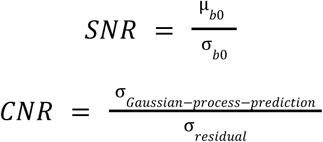

The average SNR/CNR within each map was then computed as the average within a brain mask. To improve comparability across broader datasets with different spatial resolution and scan lengths, for the DWI data we additionally report SNR_effective_, CNR_effective_ metrics normalized for differences in voxel size and the number of volumes^109^:

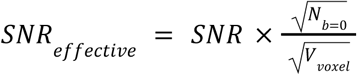

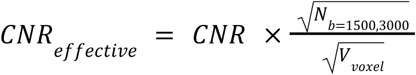

Additional QC measures included registration performance indexed by the ‘mincost’ value returned by Boundary-Based Registration (BBR)^111^, as well as ICA-derived component summaries (noise, signal, and total component counts) before and after the Reclean reclassification step (see Supplementary Fig. S2). Notably, the mincost values are useful to identify outliers within a consistently acquired dataset, but are not normalized for comparison across datasets. The ICA component summaries provide an overview of the total number of components (which tends to be higher in noisier data) and the ratio of noise-labeled components to the total number of components. These QC outputs will be shared alongside the harmonized IDPs to support transparency and secondary analyses.

### Imaging Derived Phenotype (IDP) extraction

To support broad reuse of the HARMONY data resource, we extracted a set of IDP families spanning structural, functional, and diffusion modalities. Since neuroimaging features can range from low-dimensional to very high-dimensional representations (d), we intentionally included both lower- and higher-dimensional atlases to enable dimensionality comparisons. This design supports evaluation of how feature dimensionality influences model stability, interpretability, and generalizability.

#### Structural MRI IDPs

For structural MRI, we extracted T1w-derived regional volumes, cortical thickness, and T1w/T2w ratio derived cortical myelin^55,103^. Cortical and subcortical morphology measures were derived from the HCP structural pipelines and summarized using standard FreeSurfer parcellations (DKT for cortical (d=64) and ASEG for subcortical regions (d=66)), and using the higher resolution HCP-MMP1.0 (HCP2016/Glasser) atlas (d=360) to enable dimensionality comparisons. Myelin-related contrast was quantified using the HCP-style T1w/T2w myelin map and means were extracted for these same atlases. Structural IDPs were extracted using Connectome Workbench utilities.

#### Resting state fMRI IDPs

For resting-state fMRI, we focused on measures commonly used for individual-difference and brain–behavior analyses: functional amplitude and functional connectivity. We extracted both low- and high-dimensional representations to capture complementary scales of functional organization. IDPs were derived from a 25-dimensional group ICA decomposition reflecting canonical resting state networks (Supplementary Fig. S6a) and from the higher-dimensional HCP2016FS atlas (d=379). For the 25-dimensional ICA features, we performed dual regression analysis against the 25 components derived from group-ICA performed in the Human Connectome Project Young Adult dataset. This external template set of components was used to facilitate comparisons in the literature and to avoid challenges with bias that may result from unbalanced samples in the HARMONY cohort including control-case and child-adult ratios. Subject-specific timeseries output from stage 1 of dual regression were used to calculate amplitude and connectivity IDPs as described below. For the HCP2016FS atlas, we included the 360 cortical brain areas defined in the HCP-MMP1.0 parcellation^74^ combined with 19 subcortical brain regions defined in the ASEG atlas to ensure whole-brain representation.

Functional amplitudes were calculated by taking the standard deviation of the extracted timeseries for each IC network and for each HCP2016FS parcel respectively. As such, functional amplitudes reflect the overall strength of signal fluctuations in resting state data indicative of regional synchrony^112^. For amplitude measures, this yielded 25 and 379 features, respectively.

For connectivity, we extracted both full and partial connectivity. Full correlation connectivity was defined as the Pearson’s correlation between pairwise timeseries of IC networks (300 unique edges) or HCP2016FS parcels (71,631 unique edges) respectively. Partial correlation estimates pairwise connectivity after controlling for the timeseries of all other networks/parcels. Partial (correlation) functional connectivity was calculated by taking the inverse of the covariance matrix and applying L1 regularization. Regularization strength for partial connectivity estimation was optimized to avoid over-regularization and preserve meaningful between-subject variability. Optimization was performed as a grid-search across lambdas ([0.01 0.05 0.1 0.5 1 2 3 4 5 10]), and evaluated based on the correlation between each subjects’ partial connectivity matrix at that lambda and the group-averaged partial connectivity matrix (with minimal regularization; λ=0.01). The optimal lambda was chosen as the lowest possible regularization value at which the median (across subjects) for this correlation passed r=0.5. The optimal lambda values scaled with dimensionality (λ=0.01 ICA-25 IDPs; λ=1 for ICA-200 IDPs; λ=0.5 for bilaterally collapsed HCP2016FS IDPs [see below]; λ=2 for HCP2016FS IDPs; Supplementary Fig. S7). Importantly, homologous regions across left and right hemispheres often show very high covariation, which risks removal of too much signal when using partial connectivity. Therefore, for the HCP2016FS parcellation specifically, which has a high degree of left-right homology, we also generated a bilaterally collapsed version of the HCP2016FS parcellation by averaging left–right homologous parcels, producing a 199-node representation (yielding 19,701 IDPs for unique edges). Functional connectivity IDPs were computed using FSLNets. Downstream analysis used only the upper triangle of the connectivity matrix (excluding the diagonal) to avoid redundant entries, and applied Fisher’s r-to-z transformation.

#### Diffusion IDPs

For diffusion MRI, while HCP-style acquisitions typically include multiple shells (e.g., b=0, 1500, and 3000 s/mm² in the case of the HARMONY cohorts), our primary diffusion IDPs were fractional anisotropy (FA) and mean diffusivity (MD) estimated with FSL DTIFIT^113^, which fits a single diffusion tensor under a linear Gaussian model assumption^114^. Because this tensor model is most appropriate for lower b-values and can be biased at higher b-values, we excluded the b=3000 shell prior to tensor fitting and computed FA/MD using the remaining b=1,500 (and b=0) diffusion volumes^115^.

For neurite-related white matter microstructure IDPs, we used a computationally efficient implementation of Neurite Orientation Dispersion and Density Imaging (NODDI) for all multiple shells (b=0, 1500, 3000). Specifically, we performed the NODDI-Watson^116^ implementation available in the Accelerated Microstructure Imaging via Convex Optimization (AMICO) framework^117^. From this, we derived IDP types for neurite density index (NDI), orientation dispersion index (ODI) and isotropic volume fraction (IVF).

For both DTIFIT and NODDI-derived results, IDPs were calculated for the Johns Hopkins University White-Matter Labeled (JHU81) tracts. We additionally generated skeletonized diffusion IDPs using tract-based spatial statistics (TBSS) to obtain tract-centered measures. For feature extraction, we first constructed a HARMONY-specific TBSS skeleton from 839 HARMONY participants sampled across all four sites (Supplementary Fig. S6b). Participants with limited diffusion field-of-view coverage (coverage truncated above the middle cerebellar peduncle or above the pontomedullary junction), were excluded from skeleton construction as such scans could compromise accurate estimation of the FA-derived skeleton. The resulting skeleton mask was then applied to all subjects.

### Imaging Derived Phenotype (IDP) harmonization

To mitigate undesirable site-related effects, we harmonized imaging-derived phenotypes (IDPs) across cohorts using ComBat-GAM^63^ (fit on controls) and CovBat-GAM^77^, with age and sex included as covariates in all models. Age was modeled as a nonlinear smooth term using k=5 for GAM, whereas sex was modeled as an additional categorical covariate. A particular challenge in HARMONY was that symptom severity differed systematically across sites (see Supplementary Fig. S1). Because this variance is biologically relevant and central to downstream analyses, harmonization needed to reduce site effects without removing clinically meaningful between-subject differences that were partially collinear with site.

Since HAMD was not available in the adolescent cohort (HCP-BANDA), we necessarily needed two different analysis samples, since the associated symptom was included as a covariate during the harmonization modeling to preserve variance related to that symptom. Specifically, prediction of anhedonia (SHAPS) used all four cohorts, while prediction of depression severity (HAMD) necessarily excluded HCP-BANDA.

We considered two strategies to preserve depression severity effects during harmonization. The first was to include symptom severity as an additional covariate in the harmonization model, alongside age and sex. The second was to estimate harmonization parameters using only healthy controls and then apply these parameters to cases. To prioritize robust implementation and future usability of the HARMONY resource, we used established methods available in the https://github.com/andy1764/ComBatFamily. At the time of analysis, the control-trained approach was available for ComBat-GAM but not for CovBat-GAM. Accordingly, ComBat-GAM was fit in controls and then applied to cases, whereas CovBat-GAM was fit using all subjects. We did not include ComBat-GAM trained on all subjects because the primary goal of this contribution was to introduce the HARMONY resource, such that an extensive comparison of harmonization techniques was beyond the scope of this work. The only severity-based covariate available for all four cohorts was SHAPS. Therefore, we used SHAPS as a covariate to preserve psychopathology in CovBat-GAM in addition to age and sex. One limitation of the use of SHAPS for this purpose is that it focuses on anhedonia and does not capture other core symptoms of anxious misery.

We evaluated harmonization performance using two complementary criteria: i) predictability of site following harmonization and ii) predictability of affective symptoms from IDPs (see below). To assess residual site effects, we performed cross-validated support vector machine (SVM) classification of study sites separately for each IDP file and harmonization approach, including unharmonized, CovBat-GAM, and ComBat-GAM (fit on controls). Cross-validation involved 100 repeats of 5-fold cross-validation, and each repetition was averaged across the 5-folds to promote a more stable output value from each repetition. Age and sex regression was also performed in each cross-validation split through a linear regression in the training data and then applying the training-derived beta weights to the test data. Residuals were used in subsequent analyses. To ensure fair comparison across IDPs with different feature dimensions, each IDP set was reduced using principal component analysis (PCA) to k=25 components, selected to match the lowest-dimensional representation (ICA25). PCA reduction was performed within each cross-validation fold to avoid leakage by fitting the PCA on training data and applying PCA scores to the test data. Classification performance was quantified using balanced accuracy, computed for each separate repetition as the average across each of the 5-folds. Site classification was performed separately in controls and cases across the four cohorts (chance = 25% for balanced accuracy).

### Multivariate brain-anxious misery association mapping

To assess brain–behavior prediction, we performed principal component regression (PCR) separately for each IDP file and separately for unharmonized, ComBat-GAM-harmonized, and CovBat-GAM-harmonized IDPs. Separate analyses were performed to predict anhedonia (SHAPS) and depression severity (HAMD). Depression severity predictions only included data from three adult sites due to unavailability of HAMD in HCP-BANDA, while CovBat-GAM predictions used harmonization that leveraged SHAPS as the covariate. Anhedonia predictions included data from all four sites. Cross-validation involved 100 repeated 5-fold shuffle splits same as site prediction. Each IDP set was also reduced to 25 principal components by PCA. Age and sex were regressed out at IDP level and to affective symptoms to correct for age and sex. PCA reduction and age-sex correction were performed within each cross-validation fold to avoid leakage. Model performance was quantified using cross-validated coefficient of determination (R^2^) as an effect-size metric. Note that coefficient of determination allows for negative R², which indicates that the model performed worse than simply using the sample mean as the prediction. Statistical significance of each R^2^ estimate was calculated by using a corrected resampled Student’s t-test to account for the dependence among repeated cross-validation estimates^118,119^. For 100 total (*n*) repeats of 5-fold cross-validation, we compared the R^2^ (d_i_) against zero as follows:

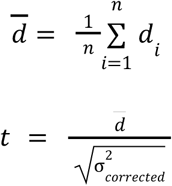

Where the corrected variance 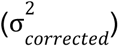 is calculated as follows:

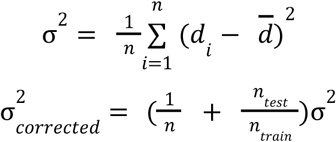

where n = 100 and n_test_/n_train_ = ¼. Given the relatively Gaussian distribution of R^2^, the one-sided p-value was subsequently obtained from the standard t-distribution with n−1 degrees of freedom:

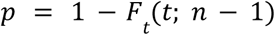

### Univariate brain-anxious misery association mapping

In addition to multivariate prediction, we performed univariate brain–behavior association analyses controlled for age and sex to provide interpretable reference results for future users of the HARMONY data resource. We focused on anhedonia, measured by SHAPS, and depression severity, measured by HAMD. Both were separately mapped to three representative IDPs covering structural, functional, and diffusion MRI: cortical–subcortical volume (HCP2016FS), partial functional connectivity (ICA25), and fractional anisotropy (skeleton JHU81). Univariate analyses were performed using ComBat-GAM-harmonized (fit on controls) IDPs only.

For each feature within each IDP file, we computed the Pearson correlation with HAMD and SHAPS across subjects after regressing age and sex with complete imaging and symptom data. This yielded feature-wise effect estimates (correlations) across all atlas regions or connectivity edges within each modality. To control for multiple comparisons, Benjamini-Hochberg false discovery rate (FDR) correction was applied separately within each IDP across all tested features. Corrected results (q < 0.05) were used to identify significant associations, and the resulting effect estimates were visualized in atlas space to illustrate the spatial distribution of univariate brain–behavior relationships. These analyses were intended to provide descriptive, interpretable benchmarks alongside the multivariate models.

## Code Availability

Updated HCP pipeline preprocessing code and information are available here: https://github.com/PersonomicsLab/HARMONY/tree/add-harmony-scripts-slurm/scripts/hcp_pipeline. All other analysis code for this article is publicly available at: https://github.com/PersonomicsLab/HARMONY/tree/add-harmony-scripts-slurm/scripts.

## Data Availability

The HARMONY data resource is available on NDA (link to be added), and on BALSA (link to be added). Raw (unprocessed) data were also shared with the NDA and are available on their website for most cohorts (https://nda.nih.gov/general-query.html?q=query=featured-datasets:Connectomes%20Related%20to%20Human%20Disease).

