## Supplementary Materials for "HARMONY: A large-scale harmonized neuroimaging dataset for research on anxious misery disorders"

#### Supplementary Information: Longitudinal design

HCP-BANDA performed two follow-up sessions repeating baseline clinical assessments, respectively 6 months (online) and 12 months (in person) after the imaging session. HCP-DES included four follow-up sessions every 3 months after the imaging session. HCP-DAM performed an in-person follow-up session 6 months after the imaging session that included clinical and functional imaging, and a telephone session 12 months after the first imaging session. On the other hand, HCP-PDC patients were recruited into three intervention cohorts (electroconvulsive therapy [ECT], ketamine, and total sleep deprivation). The longitudinal follow-up design in HCP-PDC included a full post-intervention repeat of the clinical and neuroimaging protocol, with pre-to-post time windows that varied as a function of the treatment arm. Although the HCP-PDC secondary analyses presented here focused on pre-treatment data, post-treatment neuroimaging data were also preprocessed and included in the HARMONY data resource. The ECT arm primary follow-up took place 3-4 weeks after treatment initiation, and ECT patients performed a final study visit when possible either three months after ECT treatment ended or at symptom recurrence (whichever happened first). The ketamine arm primary follow-up took place 24 hours after patients completed four serial ketamine infusions (subanesthetic dose (0.5 mg/kg) of ketamine diluted in 60cc saline delivered intravenously via pump over 40 min; 2-3 per week). A subset of ketamine patients also completed neuroimaging and clinical assessments 24 hours after their first infusion, and five weeks after ketamine treatment ended or at symptom recurrence (whichever happened first). In the total sleep deprivation arm, the primary follow-up took place directly after overnight sleep deprivation (23-25 hours after the pre-treatment session). Sleep deprivation patients also completed clinical measures (but no neuroimaging) again one week after treatment.

**Supplementary Materials**

**Table S1: HARMONY cohort characteristics**

| | Control | | | | Case | | | |
|---|---|---|---|---|---|---|---|---|
| Characteristics | BANDA | DES | DAM | PDC | BANDA | DES | DAM | PDC* |
| **Sample size** | 63 | 56 | 49 | 52 | 152 | 222 | 197 | 172 |
| **Age** (SD) | 15.2 (0.8) | 26.9 (5.3) | 27.9 (6.2) | 32.5 (11.8) | 15.5 (0.8) | 26.3 (4.9) | 28.5 (7.8) | 37.8 (12.7) |
| **Sex** | | | | | | | | |
| Female | 54.0% | 55.4% | 69.4% | 57.7% | 70.4% | 62.6% | 68.0% | 55.8% |
| Male | 46.0% | 44.6% | 30.6% | 42.3% | 29.6% | 36.5% | 32.0% | 44.2% |
| Not Reported | 0.0% | 0.0% | 0.0% | 0.0% | 0.0% | 0.9% | 0.0% | 0.0% |
| **Race** | | | | | | | | |
| White | 76.2% | 57.1% | 61.2% | 40.4% | 78.9% | 42.3% | 63.5% | 68.0% |
| Asian | 6.3% | 33.9% | 12.2% | 19.2% | 2.6% | 35.1% | 9.6% | 9.3% |
| Black or African American | 3.2% | 1.8% | 22.4% | 21.2% | 2.6% | 3.2% | 20.8% | 5.8% |
| More than one race | 12.7% | 1.8% | 0.0% | 7.7% | 14.5% | 10.4% | 1.5% | 7.0% |
| Unknown or not reported | 1.6% | 5.4% | 4.1% | 5.8% | 0.7% | 7.7% | 4.6% | 4.1% |
| Other Non-White | 0.0% | 0.0% | 0.0% | 5.8% | 0.0% | 0.0% | 0.0% | 5.2% |
| American Indian/ Alaska Native | 0.0% | 0.0% | 0.0% | 0.0% | 0.0% | 1.4% | 0.0% | 0.6% |
| Hawaiian or Pacific Islander | 0.0% | 0.0% | 0.0% | 0.0% | 0.7% | 0.0% | 0.0% | 0.0% |
| Clinical information | | | | | | | | |
| **Depression Severity (HAMD)** | - | 4.64 (5.98) | 0.92 (1.67) | 1.08 (1.68) | - | 10.85 (6.41) | 13.49 (6.03) | 18.66 (5.56) |
| **Anhedonia (SHAPS)** | 1.38 (2.53) | 7.33 (8.87) | 4.55 (5.10) | 7.16 (7.26) | 2.84 (2.90) | 12.96 (7.65) | 14.67 (7.79) | 21.98 (7.81) |
| **Baseline use of antidepressant** (% of sample size) | 0.0% | 0.0% | 0.0% | 0.0% | 0.0% | 6.3% | 18.3% | 53.9% |
| **Antidepressant classes** (% of user) | - | - | - | - | - | | | |
| SSRIs | | | | | | 92.9% | 66.7% | 47.6% |
| NDRIs | | | | | | 14.3% | 22.2% | 47.6% |
| SNRIs | | | | | | 0.0% | 16.7% | 36.9% |
| SMS/SARIs | | | | | | 7.1% | 18.9% | 24.3% |
| MAOIs | | | | | | 0.0% | 2.8% | 2.9% |
| Others | | | | | | 0.0% | 0.0% | 10.7% |

*Antidepressants included SSRIs: Selective Serotonin Reuptake Inhibitors, NDRIs: Norepinephrine and Dopamine Reuptake Inhibitors, SNRIs: Serotonin-Norepinephrine Reuptake Inhibitors, SMS: Serotonin Modulator and Stimulator, SARIs: Serotonin antagonist and reuptake inhibitors, MAOIs: Monoamine Oxidase Inhibitors, and Others including Tricyclic Antidepressant (TCA), Atypical Antidepressant, and Esketamine.*

** For HCP-PDC, psychiatric medication use among cases included any antidepressant (53.9%), lithium (3.7%), benzodiazepines (18.8%), anticonvulsants/mood stabilizers (20.4%), typical antipsychotics (0.5%), atypical antipsychotics (16.8%), stimulants (13.1%), and sleep medications (7.3%).*

### Supplementary Materials

#### Figure S1: HARMONY case characteristics

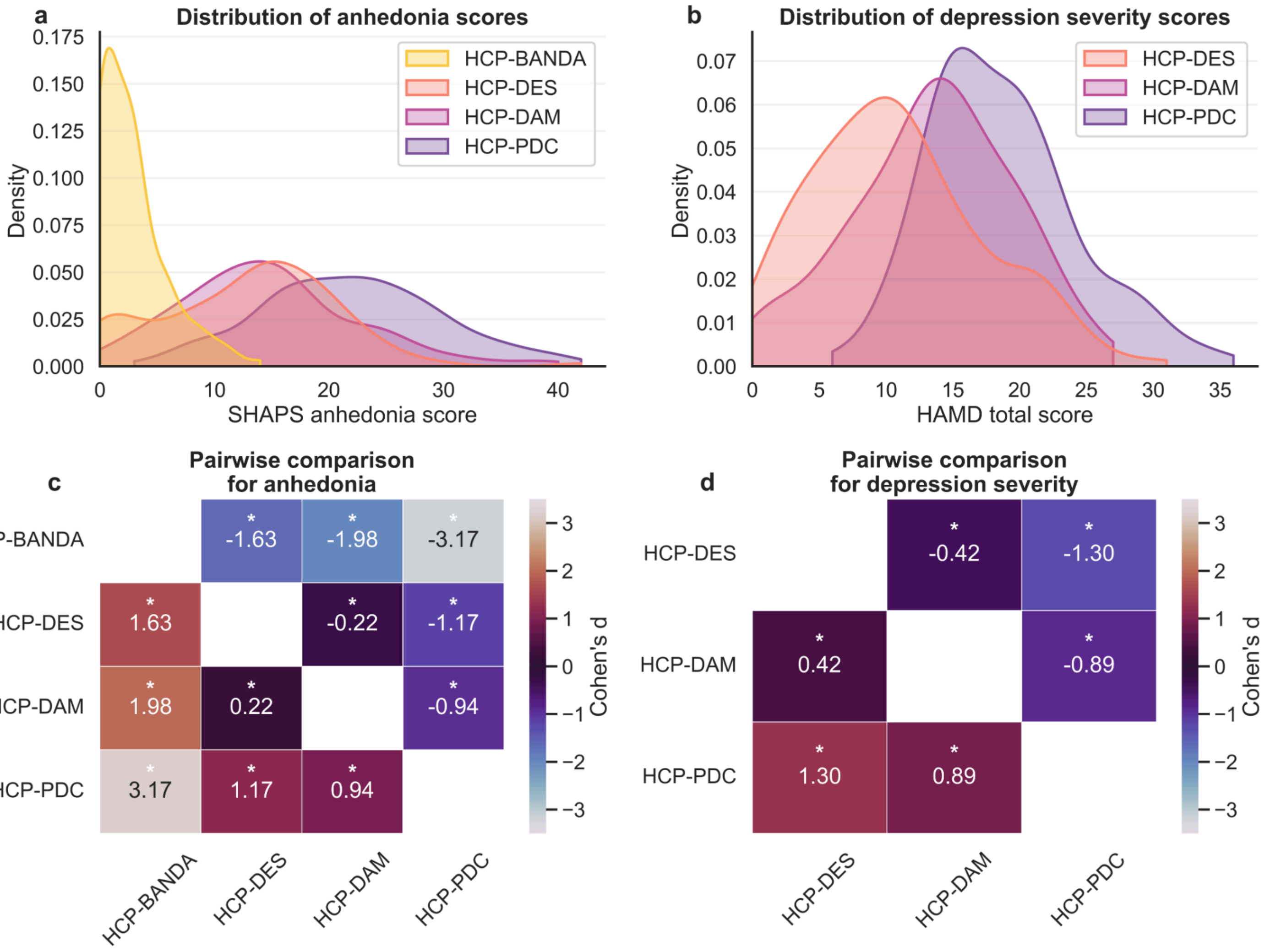

Figure S1: (a) Kernel density estimates (KDE) of baseline anhedonia (SHAPS) and (b) KDE of baseline depression severity (HAMD) for patient groups across the cohorts. Pairwise comparisons between cohorts visualized as heatmaps of Cohen's d effect sizes for (c) anhedonia and (d) depression severity. Each cell represents the standardized mean difference between the row and column cohorts. Asterisks denote statistically significant differences ($p < 0.05$, Welch's t-test). HAMD was not collected in HCP-BANDA.

**Supplementary Materials**

**Table S2: Full acquisition parameters (extended from Tozzi et al., 2021)**

| Parameter | HCP-BANDA | HCP-DAM | HCP-PDC | HCP-DES |
|---|---|---|---|---|
| Scanner | Siemens Prisma 3T | Siemens Prisma 3T | Siemens Prisma 3T | GE Discovery MR750 3T or UHP 3T[a] |
| Coil | 64-channel Siemens HeadNeck (52 "head" elements used) | 64-channel Siemens HeadNeck (52 "head" elements used) | 32-channel Siemens | 32-channel Nova head coil |
| Max. gradient strength (mT/m) | 80 | 80 | 80 | 50 |
| T1-weighted imaging (T1w) | | | | |
| Resolution ($mm^3$) | .8 × .8 × .8 | .8 × .8 × .8 | .8 × .8 × .8 | .8 × .8 × .8[b] |
| FoV ($mm^3$) | 256 × 240 × 167 | 256 × 240 × 167 | 256 × 240 × 167 | 256 × 256 × 184 |
| Slice oversampling (%) | 23.1 | 23.1 | 7.7 | 0 |
| TR (ms) | 2400 | 2400 | 2500 | 2840, or 3000 |
| TE (ms) | 2.18 | 2.22 | 1.81/3.6/5.39/7.18 | 3.55 |
| TI (ms) | 1040 | 1000 | 1000 | 1060 |
| Flip angle | 8 | 8 | 8 | 8 |
| Fat suppression | water excitation | water excitation | water excitation | not applied |
| Bandwidth (Hz/px) | 220 | 220 | 740 | 195 |
| Parallel imaging | 2 | 2 | 2 | 2 × 1.25 |
| Max. vNav reacquisition | 24 (~60 s) | Not used | 30 (~75 s) | PROMO (Max rescan=300 s) |
| T2-weighted imaging (T2w) | | | | |
| Resolution ($mm^3$) | .8 × .8 × .8 | .8 × .8 × .8 | .8 × .8 × .8 | .8 × .8 × .8[b] |
| FoV ($mm^3$) | 256 × 240 × 167 | 256 × 240 × 167 | 256 × 240 × 167 | 256 × 256 × 184 |
| Slice oversampling (%) | 0 | 0 | 7.7 | 0 |
| TR (ms) | 3200 | 3200 | 3200 | ~2500-3000 (subject specific) |
| TE (ms)[c] | 564 | 564 | 564 | ~75-80 (subject specific) |
| Number of Echoes | 303 | 314 | 303 | 120 |
| Echo train length (ms) | 1166 | 1102 | 1166 | |
| Bandwidth (Hz/px) | 744 | 744 | 744 | 781 |
| Parallel imaging | 2 | 2 | 2 | 1.9 × 1.9 |

**Supplementary Materials**

| | | | | |
|---|---|---|---|---|
| Max. vNav reacquisition | 18 (~60 s) | Not used | 25 (~80 s) | PROMO (Max rescan=300 s) |
| Diffusion magnetic resonance imaging (dMRI) | | | | |
| Resolution ($mm^3$) | 1.5 × 1.5 × 1.5 | 1.5 × 1.5 × 1.5 | 1.5 × 1.5 × 1.5 | 1.5 × 1.5 × 1.5 |
| FoV ($mm^3$) | 210 x 210 x 138 | 210 x 210 x 138 | 210 x 210 x 84 | 210 x 210 x 138 |
| b-values ($s/mm^2$) | 1500/3000 | 1500/3000 | 1500/3000 | 1500/3000 |
| Diffusion directions per shell | 93 (+7 b0) | 93 (+7 b0) | 93 (+7 b0) | 76 (+6 b0) |
| PE direction | AP/PA | AP/PA | AP/PA | AP/PA |
| Number of runs | 4 | 4 | 4 | 4 |
| Effective echo spacing (ms) | 0.69 | 0.69 | 0.69 | 0.81 |
| Bandwidth (Hz/px) | 1700 | 1700 | 1700 | 1785 or 3571 |
| Partial Fourier factor | 0.75 | 0.75 | 0.75 | 0.61 |
| Multiband factor | 4 | 4 | 4 | 4 |
| TR (ms) | 3230 | 3230 | 3230 | 3200 or 3335 |
| TE (ms) | 89.2 | 89.2 | 89.2 | 82.3 or 86.8 |
| Total scan time | 21 min 32 s | 21 min 32 s | 21 min 32 s | 17 min 30 s |
| Functional magnetic resonance imaging (fMRI) | | | | |
| Resolution ($mm^3$) | 2 × 2 × 2 | 2 × 2 × 2 | 2 × 2 × 2 | 2.4 × 2.4 × 2.4, or 2.6 x 2.6 x 2.6 |
| FoV ($mm^3$) | 208 × 208 × 144 | 208 × 208 × 144 | 208 × 208 × 144 | 220.8 × 220.8 × 144 |
| Number of frames (after discarding the first 10 frames) | 410 | 410 | 478 | 423 |
| PE direction | AP/PA | AP/PA | AP/PA | AP/PA |
| Number of runs | 4 | 4 | 2 | 4 |
| Effective echo spacing (ms) | 0.58 | 0.58 | 0.58 | 0.52 - 0.55 |
| Bandwidth (Hz/px) | 2290 | 2290 | 2290 | 5435, 5556, or 6098 |
| Partial Fourier factor | 1 | 1 | 1 | 1 |
| Flip angle (degree) | 52 | 52 | 52 | 54 |
| Multiband factor | 8 | 8 | 8 | 6 |
| TR (ms) | 800 | 800 | 800 | 710 |
| TE (ms) | 37 | 37 | 37 | 30 |

**Supplementary Materials**

| | | | | |
|---|---|---|---|---|
| Total scan time | 21 min 52 s | 21 min 52 s | 12 min 45 s | 20 min 1 s |

[a] The scanner for HCP-DES was upgraded over the course of the study. The site physicist worked to make the scans acquired on the UHP as similar as possible to those acquired on the MR750, including “down-rating” the gradient performance during the dMRI scans.

[b] Acquisition resolution; in a handful of sessions, data was interpolated to 0.5 x 0.5 x 0.8 mm during reconstruction at the scanner

[c] Siemens reports the TE of its T2-weighted long-echo-train, variable flip angle, fast/turbo spin-echo sequence (‘SPACE’) as the TE at the center of k-space. GE reports the TE of its analogous sequence (‘CUBE’) as the contrast-equivalent (“apparent”) TE. The “apparent” TE of the Siemens SPACE sequence is not reported when the echo train length exceeds 1000 ms, but is likely around 140 ms.

#### Supplementary Materials

#### Table S3: Surface-based registration manual QC codes

| Category | Code | QC flag (short label) | Description | Occurrence in HARMONY |
|---|---|---|---|---|
| **B. Surface imperfections (unedited)** | **B1** | White surface imperfections | Errors in white matter surface placement | HCP-BANDA: 41 (out of 207)<br>HCP-DAM: 85 (out of 246)<br>HCP-PDC: 116 (out of 226)<br>HCP-DES: 36 (out of 278) |
| | B1a | White too deep | White surface cuts into white matter too far | |
| | B1b | White too superficial | White surface sits too close to cortex/gray | |
| | B1c | Other (missed gyri/sulci) | Missed folds / topology placement errors | |
| | **B2** | Pial surface imperfections | Errors in pial (outer) surface placement | |
| | B2a | Pial too deep | Pial surface cuts into cortex too far | |
| | B2b | Pial too superficial (vessels/sinus) | Includes blood vessels/sinuses | |
| | B2c | Pial too superficial (dura) | Includes dura mater | |
| | B2d | Other (structure inclusion) | Includes non-cortical structures | |
| | **B3** | White & pial errors due to VR spaces / WM age spots | Surface errors driven by ventricles/VR spaces or WM hyperintensities/age spots | |
| **C. Myelin map quality issues** | **C1** | Choppy (T1/T2 intensity issues) | T1/T2 bias/intensity mismatch causing artifacts | HCP-BANDA: 40 (out of 207)<br>HCP-DAM: 29 (out of 246)<br>HCP-PDC: 8 (out of 226)<br>HCP-DES: 17 (out of 278) |
| | **C2** | Choppy (low scan quality) | Low SNR/poor acquisition quality | |
| | **C3** | Aging effects (atrophy) | Atrophy-related mapping artifacts | |
| | **C4** | Choppy/streaking (motion) | Motion-induced striping/choppiness | |
| | **C5** | Misidentified CeS (fixable by MSMAll) | Central sulcus mis-ID; MSMAll may correct | |
| **D. Miscellaneous issues** | **D1** | FNIRT registration issues | Nonlinear registration problems | HCP-BANDA: 15 (out of 207)<br>HCP-DAM: 5 (out of 246)<br>HCP-PDC: 1 (out of 226)<br>HCP-DES: 2 (out of 278) |
| | **D3** | aparc+aseg issues | Parcellation/segmentation label errors | |
| **E. Manually edited** | **E1** | FS wm.mgz edited | White matter segmentation manually corrected | HCP-BANDA: 0 (out of 207)<br>HCP-DAM: 11 (out of 246)<br>HCP-PDC: 10 (out of 226)<br>HCP-DES: 21 (out of 278) |
| | **E2** | FS brainmask.mgz edited | Brainmask manually corrected | |
| | **E3** | FS control points added | Added control points to improve surfaces/intensity | |
| | **E4** | FS -bigventricles used | Flag used for enlarged ventricles | |

| | | | |
|---|---|---|---|
| | **E5** | Alternative FNIRT config (HCA) | Special nonlinear config in PreFreeSurfer |
| | **E6** | Alternative small-head template (HCD) | Special linear template for small heads (5–7y) |
| | **E7** | Alternative large-head template (HCA) | Special linear template for large HCA heads |

**Supplementary Materials**

**Table S4: Comparison between control and case quality control metrics**

| Metric | Study | Number of control | Number of case | t-stat | Cohen's d | p-value | significant |
|---|---|---|---|---|---|---|---|
| **T1w CNR** | BANDA | 63 | 140 | -1.0351 | -0.1556 | 0.3027 | |
| | DES | 56 | 220 | 1.1357 | 0.1717 | 0.2593 | |
| | DAM | 49 | 197 | -1.0668 | -0.1624 | 0.2893 | |
| | PDC | 52 | 169 | -3.7339 | -0.4844 | 0.0003 | *** |
| **fMRI tSNR** | BANDA | 55 | 117 | 2.0703 | 0.3142 | 0.0404 | * |
| | DES | 53 | 208 | -0.5981 | -0.0844 | 0.5512 | |
| | DAM | 46 | 196 | 1.2664 | 0.2380 | 0.2103 | |
| | PDC | 48 | 163 | -1.1894 | -0.1906 | 0.2378 | |
| **DWI effective SNR (b=0)** | BANDA | 63 | 140 | 3.1316 | 0.4504 | 0.0021 | ** |
| | DES | 50 | 198 | -0.7068 | -0.1199 | 0.4821 | |
| | DAM | 46 | 181 | -0.2944 | -0.0537 | 0.7694 | |
| | PDC | 52 | 167 | 1.2701 | 0.1861 | 0.2071 | |
| **fMRI DVARS** | BANDA | 63 | 139 | -0.1777 | -0.0257 | 0.8592 | |
| | DES | 53 | 208 | 1.0702 | 0.1571 | 0.2875 | |
| | DAM | 48 | 197 | -0.4142 | -0.0663 | 0.6799 | |
| | PDC | 51 | 169 | -3.1662 | -0.4550 | 0.0021 | ** |

* for p between 0.05 and 0.01, ** for p between 0.01 and 0.001, and *** for p below 0.001

### Supplementary Materials

### Figure S2: HARMONY additional quality control measures

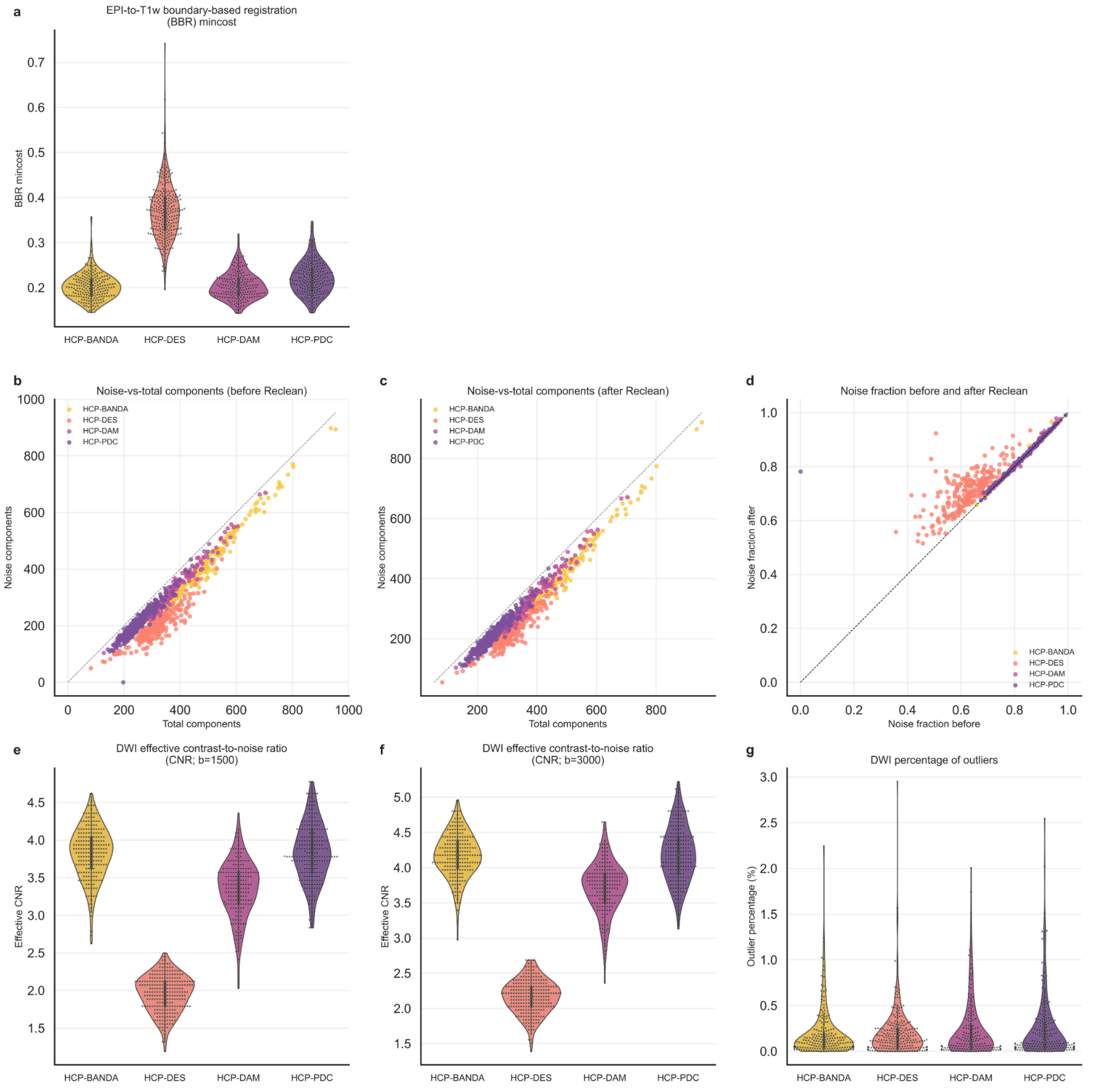

### Supplementary Materials

Figure S2: Additional quality control (QC) measures including boundary-based registration mincost (a) reflecting quality of EPI to T1w registration. A number of noise components extracted from spatial independent component (sICA) preprocessing were shown before (b) and after (c) the Reclean pipeline. The identity line shows the ceiling of possible noise components (which cannot exceed the number of total components). The noise fraction (ratio between noise and total components) before and after the Reclean pipeline (d) shows that substantially more components were reclassified from ‘signal’ to ‘noise’ in HCP-DES (collected on a GE MR750) than for the other 3 cohorts (all collected on a Siemens Prisma). This likely indicates that the original ICA-FIX classification (prior to Reclean) was just not as well-tuned for the HCP-DES data. (Note that most of the data points in panel (d) for HCP-BANDA and HCP-DAM are difficult to see, because they occur along the line-of-identity and are obscured by the overlapping data points for HCP-PDC.) Additional DWI QC measures include contrast-to-noise ratios at b=1500 s/mm$^2$ (e), and b=3000 s/mm$^2$ (f), as well as percentage of outliers (g). Results in this figure are based on both cases and controls.

### Supplementary Materials

### Figure S3: Site predictability before and after harmonization

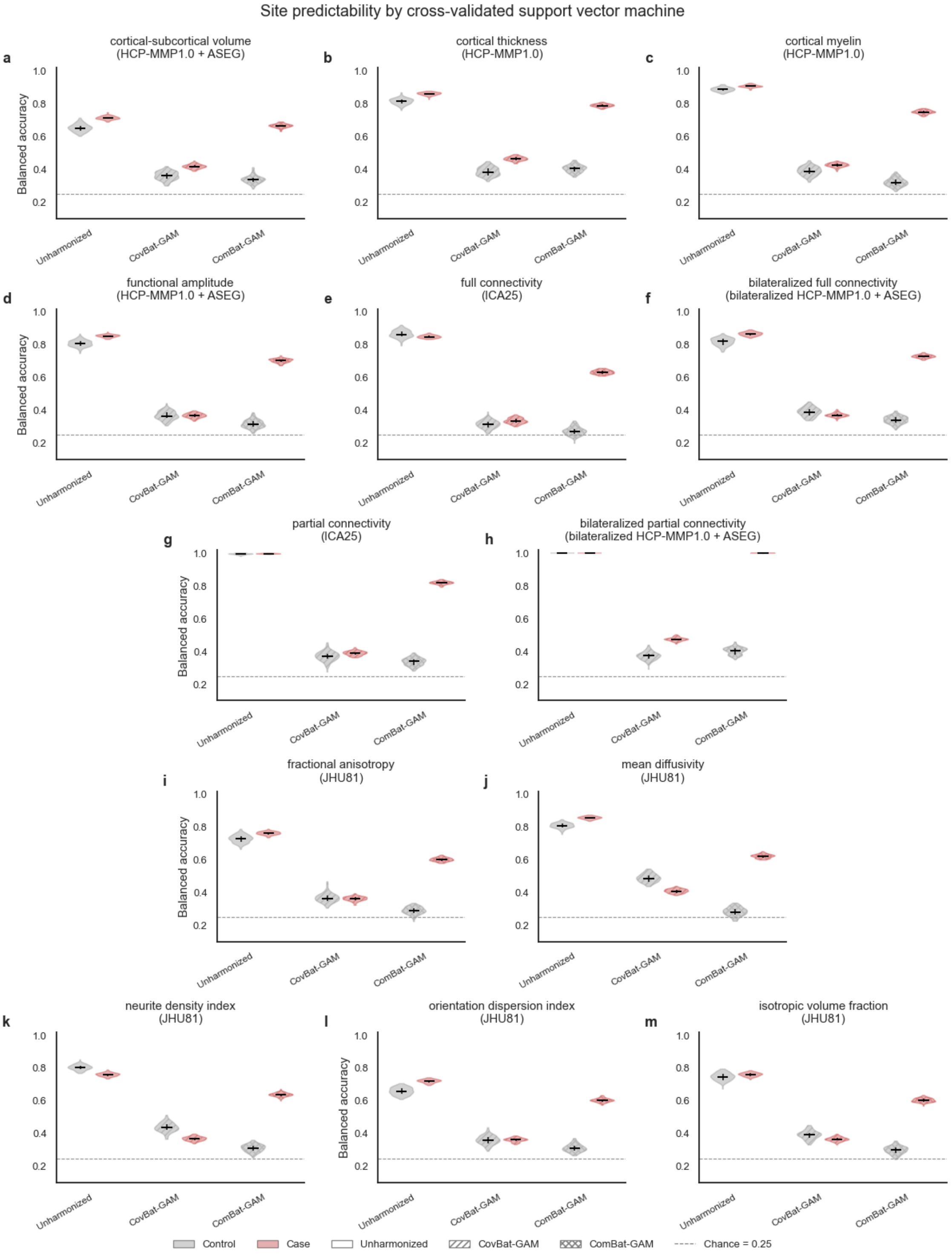

Figure S3: IDP site effects were evaluated by cross-validated SVM site classification before and after harmonization (lower accuracies reflect greater site-correction). Balanced accuracy was estimated using PCA (k=25) with 100 repetitions of 5-fold cross-validation. Results showed effective harmonization (i.e., removal of site classification accuracy) for both ComBat-GAM (fit on controls) and CovBat-GAM in controls (gray) for 13 representative IDP types: brain morphometry (a-b), myelin (c), functional amplitude (d), functional connectivity (e-h), diffusion-tensor white matter microstructure (i-j), and neurite-related white matter microstructure (k-m).

### Supplementary Materials

### Figure S4: Affective symptoms predictability after ComBat-GAM harmonization across IDPs

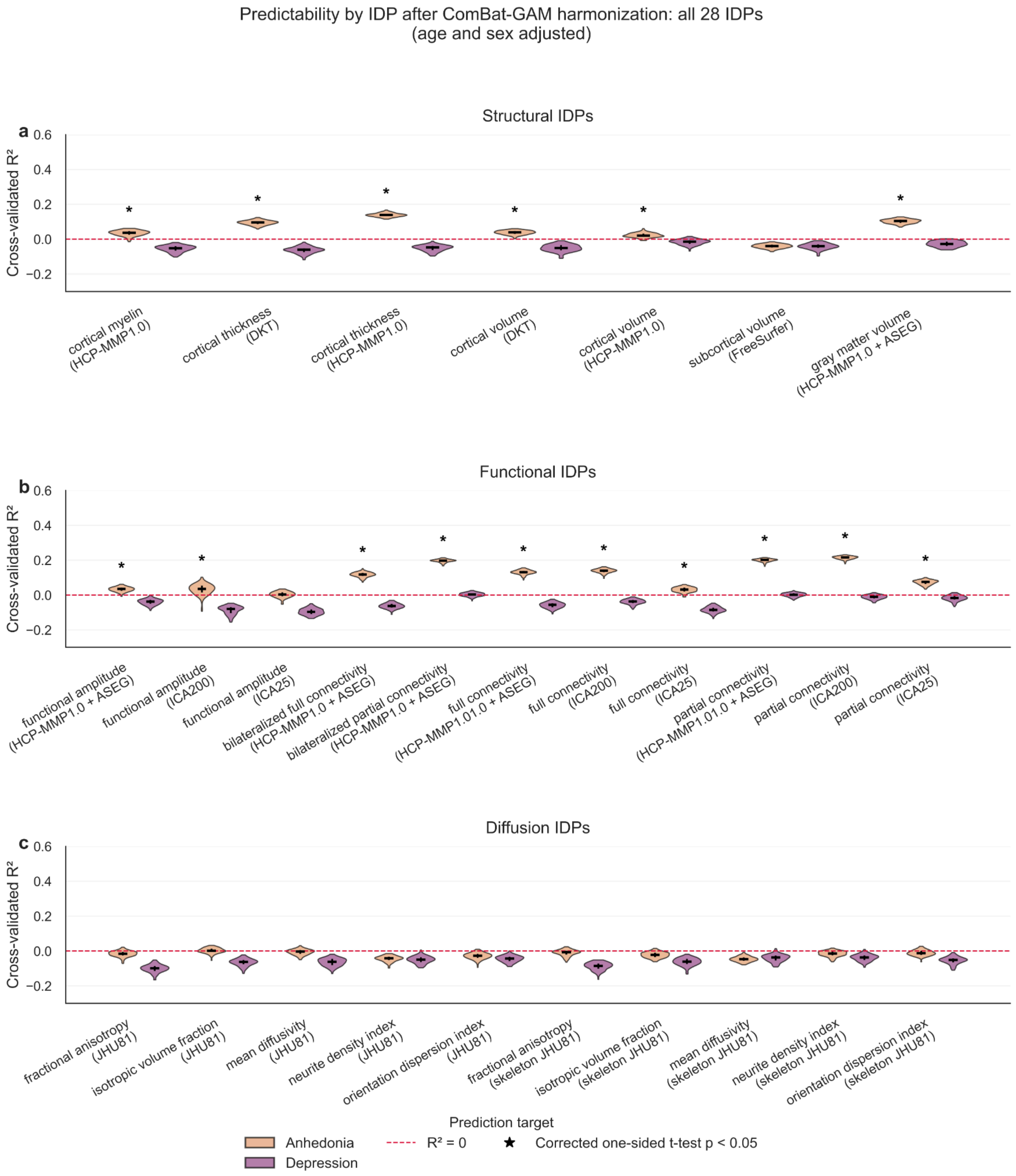

Figure S4: Multivariate brain-symptom association results for all 28 imaging derived phenotypes (IDPs) covering three imaging modalities: structural (a), functional (b), and diffusion (c) MRI. Principal component regression (k=25) with 100 repetitions of 5-fold cross-validation were performed after adjusting for age and sex. Note that $R^2$ is calculated from the sum squared error (rather than the squared correlation) and can therefore take on negative values indicative of poor model performance. Anhedonia results (orange) included all 4 cohorts and depression results (purple) included only the 3 adult cohorts because HAMD was not available in HCP-BANDA. Results in this figure are based on cases only.

**Supplementary Materials**

**Table S5: Version information for processing stages**

| HCP Pipeline | BANDA | DES | DAM | PDC |
|---|---|---|---|---|
| dcm2niix<br><br>muxarcepi2 | v1.0.20171017<br>-<br>- | v1.0.20211006<br>v1.0.20220720<br>dcc755d | v1.0.20171017<br>-<br>- | v1.0.20171017<br>-<br>- |
| Structural preprocessing | 0.96.2 | 0.100.0 | 0.96.2 | 0.96.2 |
| Structural preprocessing (manual edits) | 0.96.2 | 0.100.0 | 0.96.2 | 0.96.2 |
| Transmit bias correction | 1.3.5 | 1.3.5 | 1.3.5 | 1.3.5 |
| Functional MRI preprocessing | 0.96.2 | 0.100.0 | 0.96.2 | 0.96.2 |
| Surface alignment (MSMAll) | 0.96.2 | 0.96.2 | 0.96.2 | 0.96.2 |
| ICA-FIX denoising | 0.96.2 | 0.96.2 | 0.96.2 | 0.96.2 |
| Automated re-cleaning | 0.99.2 | 1.3.3 | 0.99.2 | 0.99.2 |
| Temporal ICA denoising | 0.99.3 | 1.3.3 | 0.99.3 | 0.99.3 |
| Diffusion MRI preprocessing | 0.96.2<br>0.100.0 | 0.100.0 | 0.96.2<br>0.100.0 | 0.96.2<br>0.100.0 |
| Diffusion modeling (BEDPOSTX) | 0.96.2 | 0.96.2 | 0.96.2 | 0.96.2 |

Values represent the QuNex version used for that particular stage of processing, except for DICOM to NIFTI conversion, for which the version/hash of the indicated converter is provided. HCP-DES data was provided to the Connectome Coordination Facility already converted to NIFTI, which is why the ‘dcm2niix’ version differs relative to the other 3 cohorts.

### Supplementary Materials

#### Figure S5: Benefits of larger sample size in HARMONY

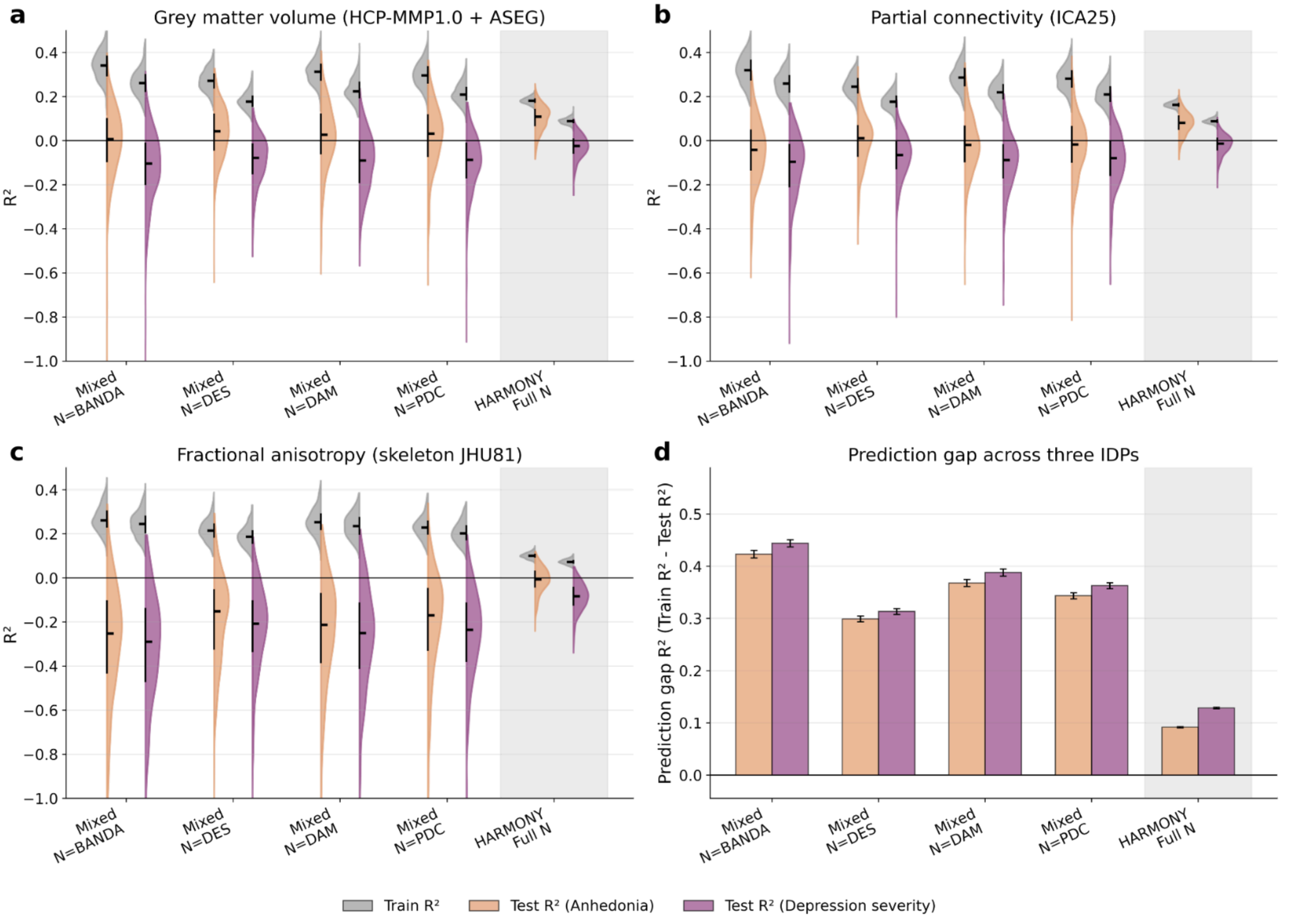

Figure S5: Benefits of larger sample size in HARMONY for three ComBat-GAM-harmonized IDPs – gray matter volume (a), partial connectivity (b) and fractional anisotropy (c). To evaluate the contribution of sample size to model prediction, we drew repeated subsamples from the pooled HARMONY dataset, focused only on cases, matched to the sample sizes of the individual cohorts and weighted by each study's proportional representation among HARMONY cases. For each IDP feature set, models were evaluated in mixed HARMONY subsamples matched to the original cohort-specific sample sizes and predicted targets availability (N=BANDA [135-137], N=DES [179-206], N=DAM [144-160], and N=PDC [167-170]) and compared with the full pooled HARMONY sample. Prediction gaps (train $R^2$ - test $R^2$) were calculated from 100 repetitions across the three selected IDPs (d). Error bars indicate standard errors of mean across 300 datapoints (3 IDPs x 100 repetitions each). The prediction gap in each of the matched-size mixed samples is comparable to the prediction gap in the original cohorts (Fig. 6), indicating that the reduction in the prediction gap in the full HARMONY cohort is due to the benefits of a larger sample size rather than cohort mixing. The results shown here for HARMONY differ slightly from those in Fig. 6, because they are based on a new set of 100 repetitions. Anhedonia and depression severity results were in orange and purple, respectively. Results in this figure are based on cases only.

### Supplementary Materials

#### Figure S6a: HCP-YA derived 25 independent components

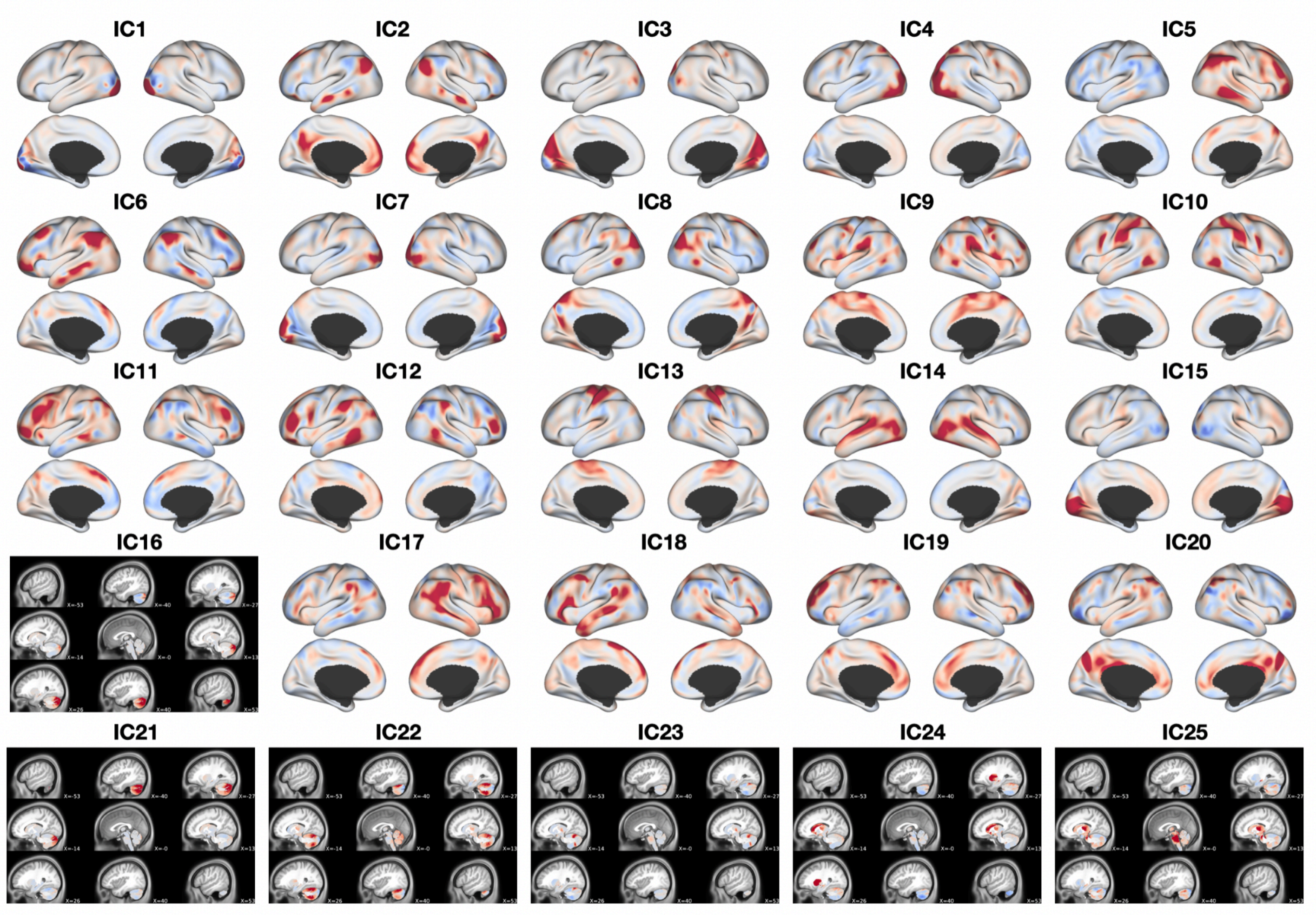

#### Figure S6b: HARMONY FA-derived skeleton mask (HCP-H869)

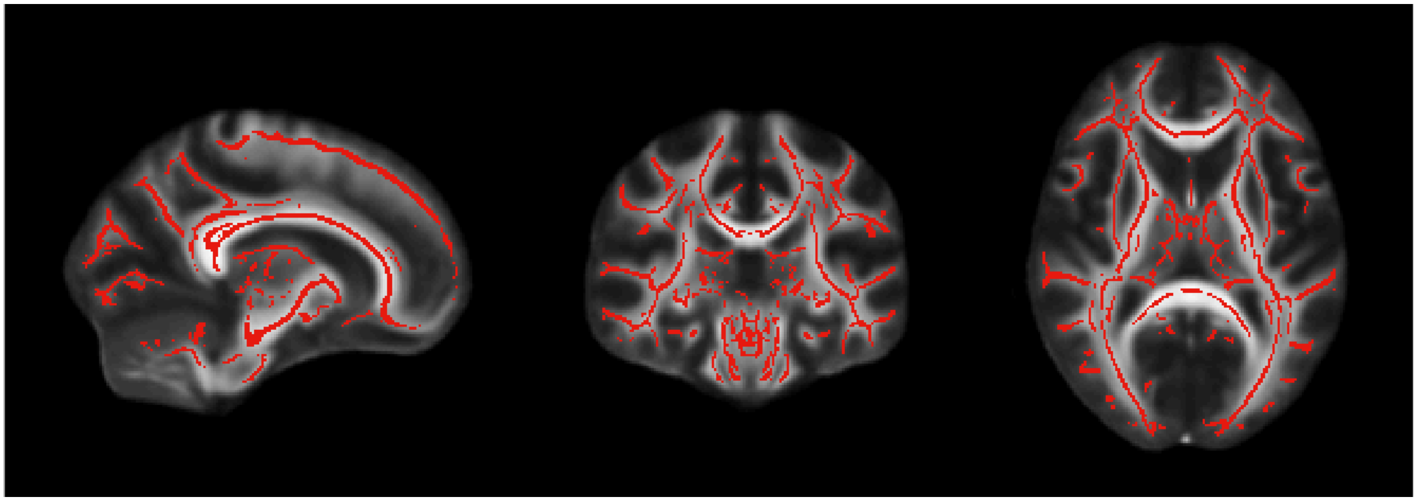

Figure S6: Group atlases used in HARMONY. Spatial independent component analysis (sICA) 25 components derived from Human Connectome Project Young Adult (Fig. 5A), and HARMONY-derived TBSS skeleton map (Fig. 5B). In panel (A), components primarily represented in the cerebellum or subcortical structures are shown on sagittal slices (rather than a surface representation).

### Supplementary Materials

### Figure S7: Regularization strength optimization for partial connectivity metrics

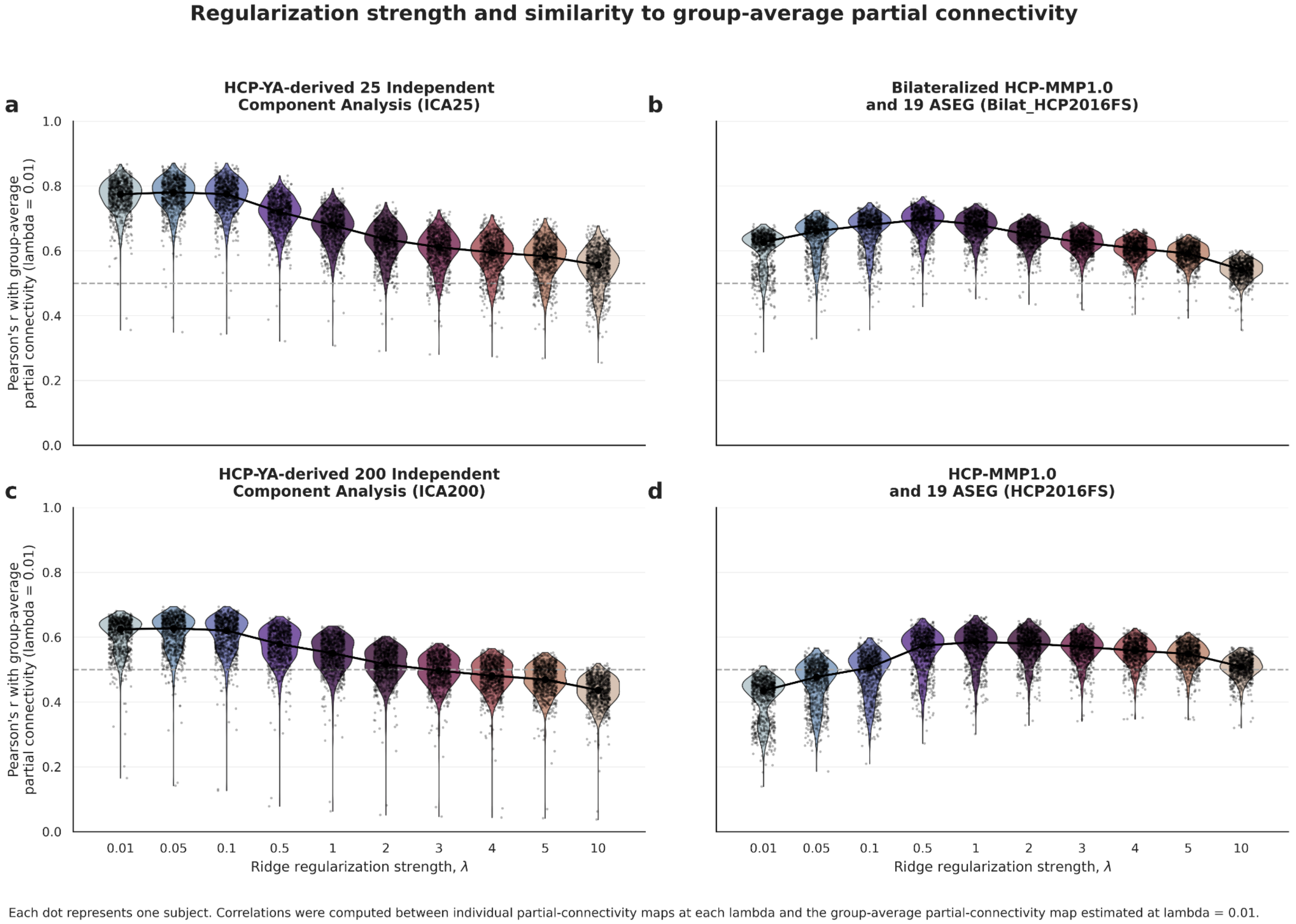

Figure S7. Grid-search optimization of regularization strength for partial connectivity estimation. Regularization strength was optimized by evaluating a range of lambda values ([0.01, 0.05, 0.1, 0.5, 1, 2, 3, 4, 5, 10]). For each lambda, partial connectivity estimates were compared with the group-average partial connectivity matrix with the smallest regularization (lambda = 0.01). The dashed gray line indicates a correlation of 0.5. The optimal lambda was selected as the highest correlation to group-average partial connectivity. This procedure identified lambda = 0.05 for ICA25 (a), lambda = 0.5 for bilateralized HCP2016FS (b), lambda = 0.05 for ICA200 (c), and lambda = 1 for HCP2016FS (d; HCP-MMP1.0 + ASEG). Results in this figure are based on both cases and controls.
